# PPML-Omics: a Privacy-Preserving federated Machine Learning method protects patients’ privacy in omic data

**DOI:** 10.1101/2022.03.23.485485

**Authors:** Juexiao Zhou, Siyuan Chen, Yulian Wu, Haoyang Li, Bin Zhang, Longxi Zhou, Yan Hu, Zihang Xiang, Zhongxiao Li, Ningning Chen, Wenkai Han, Di Wang, Xin Gao

**Author notes:** Equal contribution (Juexiao Zhou, Siyuan Chen, Yulian Wu).

## Abstract

Modern machine learning models towards various tasks with omic data analysis give rise to threats of privacy leakage of patients involved in those datasets. Despite the advances in different privacy technologies, existing methods tend to introduce too much computational cost (e.g. cryptographic methods) or noise (e.g. differential privacy), which hampers either model usefulness or accuracy in protecting privacy in biological data. Here, we proposed a secure and privacy-preserving machine learning method (PPML-Omics) by designing a decentralized version of the differential private federated learning algorithm. We applied PPML-Omics to analyze data from three sequencing technologies, and addressed the privacy concern in three major tasks of omic data, namely cancer classification with bulk RNA-seq, clustering with single-cell RNA-seq, and the integration of spatial gene expression and tumour morphology with spatial transcriptomics, under three representative deep learning models. We also examined privacy breaches in depth through privacy attack experiments and demonstrated that PPML-Omics could protect patients’ privacy. In each of these applications, PPML-Omics was able to outperform methods of comparison under the same level of privacy guarantee, demonstrating the versatility of the method in simultaneously balancing the privacy-preserving capability and utility in practical omic data analysis. Furthermore, we gave the theoretical proof of the privacy-preserving capability of PPML-Omics, suggesting the first mathematically guaranteed method with robust and generalizable empirical performance in protecting patients’ privacy in omic data.

## 1 Introduction

Individual privacy in biology and biomedicine is emerging as a big concern [1] with the development of biomedical data science in recent years. A deluge of genetic data from millions of individuals are generated from massive research projects in the past few decades, such as the Cancer Genome Atlas (TCGA) [2], the 100,000 genome project [3] and the Earth BioGenome Project (EBP) [4] from high-throughput sequencing platforms [5]. Those datasets may lead to potential leakage of genetic information and privacy concerns on ethical problems like genetic discrimination [6]. Consequently, a large amount of potentially private genetic information from modern multi-modal sequencing platforms, including bulk RNA sequencing (RNA-seq) [7], single-cell RNA sequencing (scRNA-seq) [8] and spatial transcriptomics [9] might also be exposed as more and more data are being published (**Supplementary Section 1.1**).

In addition to the data itself, another risk factor to data privacy is the wide scope of applications of machine learning, especially deep learning, which evolves rapidly by taking advantage of large datasets. A massive number of models and applications, which require training on a large and diverse dataset (either public data or in-house data), are being created, shared and applied to various areas, such as the genomics [10], medical imaging [11] and healthcare [12]. Nevertheless, individual privacy is being exposed to high risk, leading to the new concern of privacy in modern artificial intelligence (AI) [13]. As shown in **Figure 1a**, typically, sensitive data may exist in a distributed manner, where the data owners do not want to share the raw data for privacy reasons, while the aggregators want to access enough data to improve model utility. To balance the needs of both, we need a machine learning framework for distributed data that can balance utility and privacy-preserving capabilities: a secure and privacy-preserving machine learning (PPML) method.

**Fig. 1.**
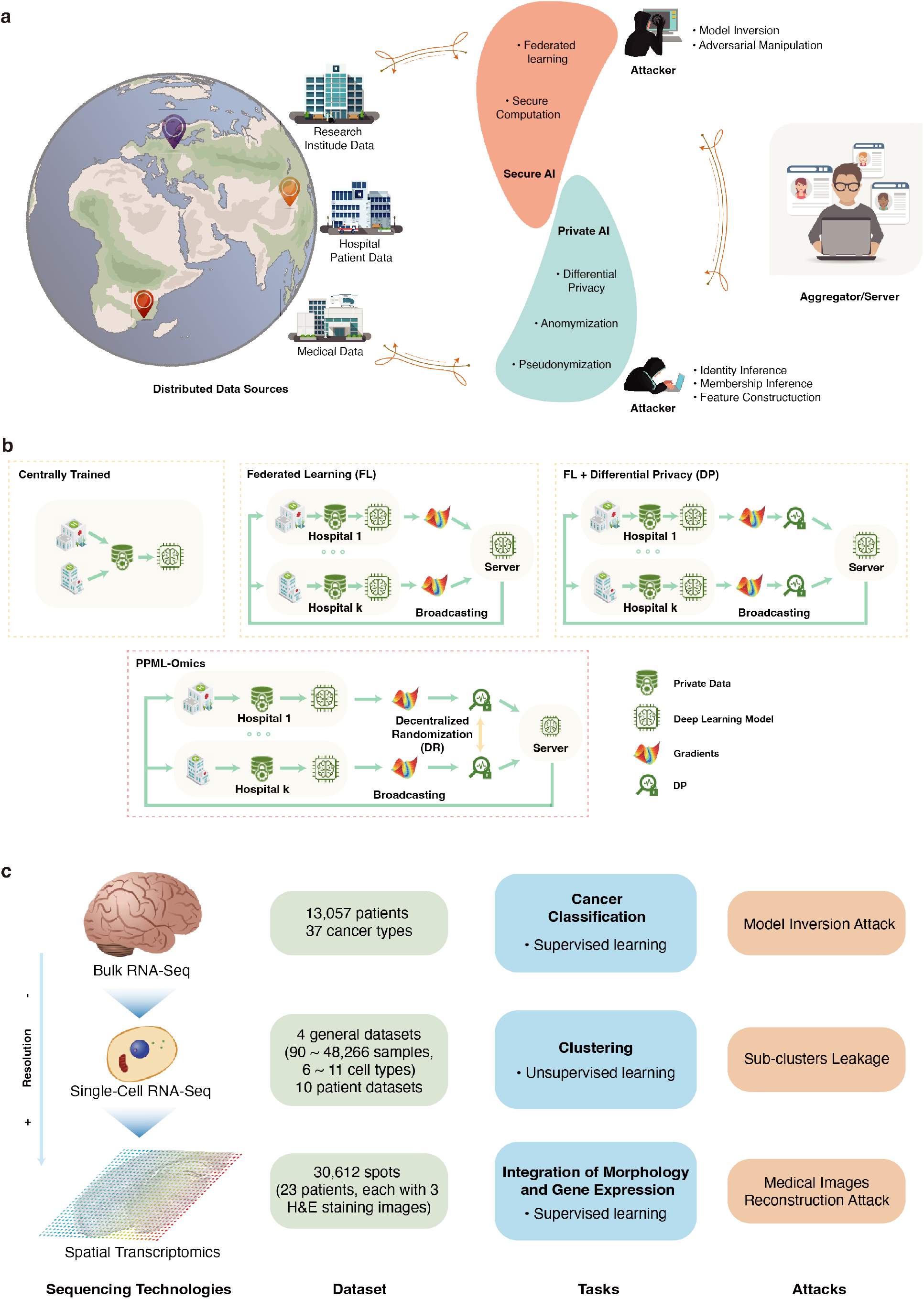
PPML-Omics: a Privacy-Preserving federated Machine Learning method protects patients’ privacy in omic data. **a**, Schematic overview of the relationships and interactions between distributed data owners, aggregator, attackers and techniques in the field of secure and private AI. **b**, Schematic overview of different methods, including centrally trained method, federated learning (FL), FL with differential privacy (DP), and PPML-Omics. **c**, Illustration of 3 representative tasks, datasets and attacks of omic data in this paper for demonstrating the utility and privacy-preserving capability of PPML-Omics, including the 1) cancer classification with bulk RNA-seq, 2) clustering with scRNA-seq and 3) integration of morphology and gene expression with spatial transcriptomics.

To alleviate the leakage of privacy, the most commonly used strategy is the anonymization or pseudonymization of sensitive data before transmitting it to the data-sharing centre [14]. Unfortunately, recent studies showed that anonymization was insufficient towards re-identification attacks [15] and linking attacks [16]. To overcome the shortness of centralized data sharing and model training, federated learning (FL) was proposed in 2017 as a data-private collaborative learning method [17]. The collaborating institutions train a machine learning (ML) model with their own data in parallel and send the model updates to the central server, which can aggregate all model updates into a consensus model without accessing the raw data. Nevertheless, the distributed nature of FL gives rise to new threats of privacy leakage caused by potentially malicious participants [18], [19], [20], [21], such as data poisoning attack [22], membership inference attack [23], [24], [25], source inference attack (SIA) [26] and data reconstruction attack [27]. Hence, exposing the trained model to a non-trusted user may also cause privacy leakage [28].

To further strengthen FL’s privacy guarantee to preserve privacy, additional privacy-enhancing modules are required. Within the most extensively studied field, multi-party computation (MPC) or multi-party homomorphic encryption (MHE) frameworks use cryptographic techniques to protect the data while enabling the training of ML models with perfect accuracy. These techniques have been used to secure FL training [29], [30], [31], [32] and achieve stronger privacy protection at the expense of computational efficiency, which might be difficult to satisfy for some clients in practice. MPC incurs a high network-communication overhead and is difficult to scale to a large number of clients, while MHE introduces high storage, computational over-heads and a single point of failure in the standard centralized setup, where one server receives all encrypted datasets to secure federated computation [33]. Besides, blockchain is also used to secure FL training such as the Swarm Learning [34]. However, applying blockchain for deep learning with FL in practice is still a challenging field due to the high communication and computing cost [35]. Meanwhile, most encryption methods still leak privacy in case of the attacker masters the corresponding decryption technique, such as the key.

Thus, an innovative and state-of-the-art solution named differential privacy (DP) was proposed as another solution [36] (**Supplementary Section 1.2**) to yield a privacy-utility trade-off, under which we could protect privacy by only sacrificing some data utility, and keep protecting privacy even if the model is stolen by an attacker. The topic of FL with DP was explored extensively in literature [37], [38], [39], [40], [41]. However, most of the aforementioned articles only reported theoretical analyses of their framework or tested on classical datasets in the field of computer science, such as MNIST and CIFAR-10, whereas only few of them applied their frameworks to real biological datasets that had more complex properties and greater intrinsic noise.

Among those works that applied FL, MPC, MHE and DP to omic data analysis, most of them utilized either the cryptographic techniques [42], [43], [44] or the DP notion [33], [42], [45], [46], [47], [48], [49] to provide formal privacy guarantees for the participants in the research of SNPs, GWAS and differential gene expression analysis [50], which are relatively narrow and specific problems in genomics studies, and whose data is obtained by post-processing the raw sequencing data (**Supplementary Section 1.3**). In addition, the methods in those articles could only be shown to be applicable to statistical solutions or traditional machine learning solutions in GWAS. In addition to SNPs, raw sequencing data saved in the matrix format and generated by high-precision and quantitative multidimensional sequencing technologies contains much more sensitive information. Apart from that, only Swarm Learning [34] discussed the application of FL with blockchain technology in the analysis of omic data. But with the notation of DP, to our knowledge, only one work discussed the application of FL-DP in cancer prediction as a solution to the competition hosted by iDASH (integrating Data for Analysis, Anonymization, SHaring) National Center for Biomedical Computing in 2020 [51]. Apart from that, there is no more work that has systematically studied and delved into the privacy protection of sequencing data from a bigger picture with the DP notation, even though raw sequencing data contains much more private information about patients than SNPs and GWAS. Meanwhile, the state-of-the-art work related to applying DP and MPC protocols in other biological tasks only reported the privacy protection in medical imaging [52], [53], which is unable to be able to generalize to omic data analysis tasks because omic data have very different characteristics to imaging data.

To find a solution that is more applicable to practical scenarios of biological problems, we proposed a robust and powerful PPML-Omics method by designing a decentralized version of the differential private federated learning algorithm (**Methods**) (**Figure 1b**). In essence, the gradients of locally trained federated machine learning models are obfuscated through differential privacy (DP) and decentralized randomization (DR) mechanisms before aggregating them at a single and non-trusted party. We applied PPML-Omics to analyze and protect privacy in real biological data from three representative omic data analysis tasks, which were solved with three different but representative deep learning models. We demonstrated how to address the privacy concern in the cancer classification from TCGA with bulk RNAseq [7], clustering with scRNA-seq [54], and the integration of spatial gene expression and tumour morphology with spatial transcriptomics [55], [56]. In addition, we examined in depth the privacy breaches that existed in all three tasks through privacy attack experiments and demonstrated that patients’ privacy could be protected through PPML-Omics as shown in **Figure 1c**. In each of these applications, we showed that PPML-Omics was able to outperform methods of comparison, demonstrating the versatility of the method in simultaneously balancing the privacy-preserving capability and utility in omic data analysis. Finally, we proved the privacy-preserving capability of PPML-Omics theoretically (**Supplementary Section 1.4**), suggesting the first mathematically guaranteed method with robust and generalizable empirical performance in the application of protecting patients’ privacy in omic data.

In summary, our contribution provides the following innovations. We introduced the DP concept and systematically studied the privacy problem of multi-omics analysis in the form of three significant application scenarios in biology. We proposed PPML-Omics to achieve a better trade-off between model performance and privacy-preserving capabilities by designing a decentralized version of the differential private federated learning with the decentralized randomization (DR) protocol based on the Fisher-Yates shuffle algorithm [57]. Additionally, we demonstrated the training of three representative deep learning models on three challenging tasks of omic data analysis using PPML-Omics and addressed the privacy concern in all three tasks, which may benefit following researchers by reminding the privacy issues in analyzing omic data. Besides, we conducted extensive experiments and showed that PPML-Omics was compatible with a wide range of omic data and biological tasks. In addition, we examined the computational performance of models trained with PPML-Omics against models trained centrally on the accumulated dataset and models trained with the FL method, FL-DP method, and FL-MHE method to demonstrate the strength of PPML-Omics under various scenarios typical in omic data analysis. Finally, we assessed the theoretical and empirical privacy guarantees of PPML-Omics and provided examples of applying state-of-the-art attacks against the models in the application of protecting patients’ privacy in omic data.

## Results

### PPML-Omics and threat models

A confederation of *N* (≥ 3) hospitals wishes to train three deep learning models for three tasks as shown in **Figure 1b,c**. Since those hospitals had neither enough data themselves nor the expertise to train models on that data, they sought the support of a model developer to coordinate the training on a central server. PPML-Omics is built on this scenario to meet the needs of federal learning training and preserve the confidentiality of the local data from attacks. In the training phase of this method, each hospital has its own private patient data as the data owner. We suppose that participants trust each other (at least semi-honest) and do not actively undermine the learning protocol. Each hospital trains a local model with its own private data and exchanges its gradients with another hospital randomly with the DR protocol before sending the updates to the server. Under this setting, each hospital is partially trusted by each other such that the gradients could be exchanged with a randomly generated partner in each epoch while the raw data cannot be accessed directly. In each epoch of training, after the DR mechanism, all hospitals may not hold their own original gradients, but rather gradients from a randomly paired hospital. Then all hospitals upload their gradients to a server that is not trusted by the hospitals as the server usually requires strong communication power, and is controlled and maintained by a third party. After integrating all gradients by the server, the server model is updated and sent to all hospitals to update all local models. Furthermore, individual participants excluding all clients are assumed as potential attackers to actively try to extract private information from other participants’ data, the transmitted gradients during the training phase of the FL method or the released pre-trained model due to curiosity. The DP-based privacy enhancement technique was introduced in PPML-Omics to prevent such behaviour. Specifically, it bounds the worst-case privacy loss for a single patient in the dataset and provides privacy guarantees to prevent model inversion/reconstruction attacks on federation participants or model owners during inference by adding noise to the gradients passed in the FL method. PPML-Omics implements DP and DR protocol to provide client-level privacy guarantees, further potentially protecting patient-level privacy. At the end of the training, all participants will hold a copy of the fully trained final model.

In Application 1, the participants try to train a model for cancer-type classification based on gene expressions from in-house bulk RNA-seq data. The server would release the final trained model and make it available to potential users. Since the released model may remember a large amount of gene expression information that is closely related to the cancer type from the patients. We assume that the attacker has auxiliary information, such as knowing that patient 1 is of cancer type A and has participated in the training of the released model. Thus, by performing the model inversion attack (MIA), the attacker could roughly know the gene expression of patient 1, resulting in a potential patient privacy breach.

In Application 2, the participants try to train a model for unsupervised clustering based on gene expressions from scRNA-seq data. The server would release the final trained model and make it available to potential users. We assume that in a real-world application scenario, users will have their own scRNA-seq data and also have our published model trained with PPML-Omics with different privacy budgets *ϵ*. The user’s treating physician needs to analyze the user’s clustering results from scRNA-seq data to determine the cell composition for medical judgment. However, the user does not want to present her clustering results to the treating physician with 100% accuracy due to privacy concerns. Since the user does not have extensive medical knowledge, the user can not modify the clustering results by herself and can only hide some of the details of the clustering results by selecting different embedding models trained with PPML-Omics with different privacy budgets *ϵ*.

In Application 3, the participants try to train a model for predicting the spatial gene expression based on highresolution images of H&E staining tissue from spatial transcriptomics data. The server would release the final trained model and make it available to potential users. Since the medical images used in the training phase contain lots of patients’ privacy, we assume that an attacker will use the improved deep leakage from gradients method (iDLG) to reconstruct the images in the training dataset by stealing the gradient transmitted between the client and the server, resulting in a potential patient privacy breach

### Application 1. Cancer classification with bulk RNA-seq

Bulk RNA-seq can directly disclose patients’ gene expression, while machine learning models trained with bulk RNA-seq can indirectly leak patients’ signature expressed genes [49]. Here, we collected data from TCGA (**Methods and Supplementary Table 1**) and compared computing resource requirements, privacy-preserving capabilities, and utility on data of different methods by working on the cancer classification task with gene expression as inputs.

To study the robustness and demonstrate the advantages of computational resources required by PPML-Omics, we used this task as a benchmark for the profiling analysis to test different methods (level 1: with or without FL, DP and DR) against varied machine learning networks (level 2) (**Figure 2a**). Depending on the differences in level 1, we tested five methods, including the centrally trained method, which was trained with one model on the entire dataset pooled on a single machine; the FL method, which was trained with separate models on the individual data owners’ subsets of the dataset following the protocol of federated learning; the FL-MHE method, in which the MHE mechanism was integrated on the FL method; the FL-DP method, in which the DP mechanism was integrated on the FL method; and PPML-Omics, in which the DR protocol was designed to achieve a better trade-off between the model performance and the privacy-preserving capability.

**Fig. 2.**
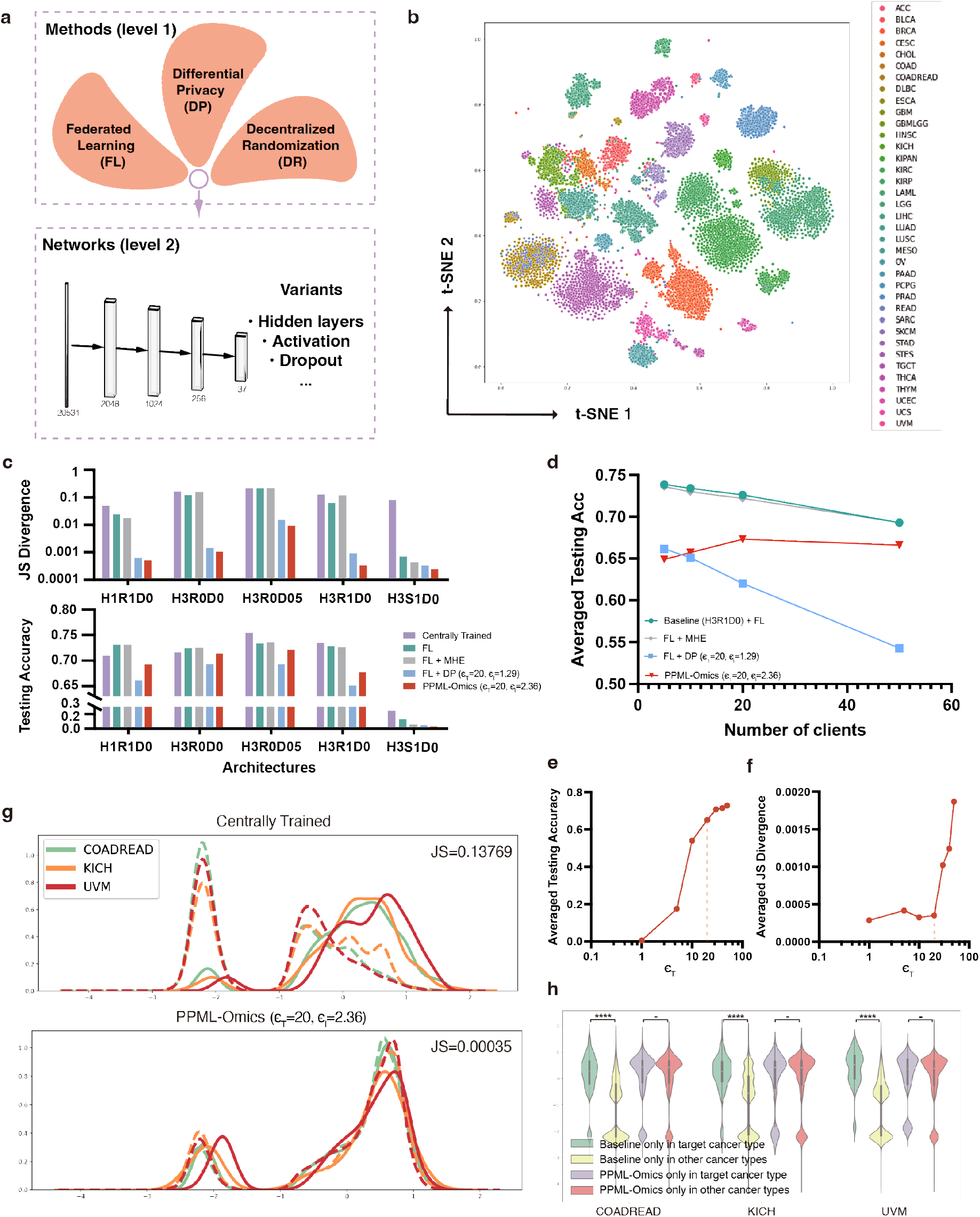
Results of cancer classification with bulk RNA-seq in Application 1. **a**, Illustration of the relationship between methods (level 1) and Networks (level 2). **b**, t-SNE plot on all patients’ data from TCGA, each data point represents one patient and colours represent cancer types. **c**, Profiling analysis of different methods, including the centrally trained method, the FL method, the FL-DP method, and PPML-Omics against different networks (level 2 in **a**) with varying numbers of hidden layers (H1: 1 hidden layer, H3: 3 hidden layers), activation function (R: ReLU, S: sigmoid), the dropout layer (D0: without dropout layer, D05: with dropout layer and *p* = 0.5). **d**, The effect of the number of clients on Application 1 of four methods. **e**, The effect of the value of *ϵ*_*T*_ on the averaged testing accuracy of PPML-Omics. **f**, The effect of the value of *ϵ*_*T*_ on the averaged JS divergence of PPML-Omics. **g**, The distribution of z-score normalized expression of reconstructed signature genes in the target cancer type (solid lines) and other cancer types (dashed lines) by the centrally trained method (baseline) and PPML-Omics from three representative cancers (COADREAD: Colorectal adenocarcinoma, KICH: Kidney Chromophobe, UVM: Uveal Melanoma). **h**, The violin plot of the distribution of z-score normalized expression of reconstructed signature genes in the target cancer type and other cancer types specifically identified by the centrally trained method and PPML-Omics. The test performed was two-sided Kolmogorov–Smirnov test, p-value annotation legend is the following: \*\*\*\**P* ≤ 0.00001, ^−^*P >*0.001. Exact P values are the following: Baseline only on COADREAD, *P* = 5.98454 *×* 10^−59^; PPML-Omics only on COADREAD, *P* = 0.03325; Baseline only on KICH, *P* = 3.59413 *×* 10^−34^; PPML-Omics only on KICH, *P* = 0.19957; Baseline only on UVM, *P* = 2.14898 *×* 10^−131^; PPML-Omics only on UVM, *P* = 0.00144.

Furthermore, for each method in level 1, variants based on the fully connected neural network (FCN) (**Figure 2a**) were tested (level 2), including the use of different numbers of hidden layers (H1: 1 hidden layer, H3: 3 hidden layers), activation functions (R: ReLU, S: sigmoid), dropout layers (D0: without dropout layer, D05: with dropout layer and p = 0.5) and values of end-to-end *ϵ*_*T*_, which can be used to further calculate the *ϵ*_*l*_ for each client in each epoch for the 37-class classification task, where the end-to-end privacy budget *ϵ*_*T*_ was a hyperparameter (**Methods**) and a smaller *ϵ*_*T*_ means that we have stricter requirements for privacy protection, resulting in the need to add more noise and lower model performance. Data visualization through t-SNE showed that the use of gene expression could effectively distinguish between different cancer types (**Figure 2b**). We trained all models to converge on the same dataset, and measured the privacy-preserving capability (**Methods**) and required computational resources, including time, RAM and GPU memory (**Supplementary Section 2.2**). Following Li’s definition of the privacy loss in [58], we used Jensen-Shannon (JS) divergence and the p-value of Kolmogorov–Smirnov (KS) test (*P*_*KS*_) between two distributions to quantify privacy leakage, where a larger JS divergence and a smaller p-value indicate a more severe privacy leakage. Compared to the centrally trained method, the FL method did not exhibit significant privacy-preserving power from the JS divergence perspective (JS divergence *>*0.1 and similar to the centrally trained method). Compared to that, the integration of the DP (FL-DP method) substantially improved the privacy-preserving power (JS divergence ≤ 0.01) as shown in **Figure 2c**. The JS divergence gradually decreased as the end-to-end privacy budget *ϵ*_*T*_ decreased (**Figure 2f**), indicating that adding higher noise (smaller *ϵ*_*T*_) allowed for stronger privacy protection. As a trade-off between privacy protection and the data utility, *ϵ* ≤ 5 is usually chosen in practice as in the handbook ‘Disclosure Avoidance for the 2020 Census: An Introduction’ [59] (**Methods**). However, there is no definite range of specific values for *ϵ* and the value of *ϵ* is dependent on the task and the data, and needs to be selected based on a comprehensive consideration of the privacy-preserving requirement and the utility. In Application 1, *ϵ*_*T*_ = 20 balanced the utility (**Figure 2e**) and the privacy-preserving capability (**Figure 2f**) of PPML-Omics. Therefore, with *ϵ*_*T*_ = 20, PPML-Omics (JS divergence=0.00035, *P*_*KS*_ = 0.52341) showed a clear advantage over the centrally trained model (JS divergence=0.13769, *P*_*KS*_ = 8.21642 × 10^−46^), the FL method (JS divergence=0.06169, *P*_*KS*_ = 1.66231 × 10^−21^) and the FL-MHE method (JS divergence=0.10567, *P*_*KS*_ = 2.01988 × 10^−39^) when we need to meet both acceptable computational requirements (**Supplementary Table 2**) and privacy protection capabilities (**Figure 2c**). Overall, PPML-Omics with *ϵ*_*T*_ = 20 could meet the need for privacy protection under the practical scenario in Application 1 and the computational resource requirements of PPML-Omics were notably user-friendly.

Adding noise to gradients with DP has an amplification effect [60], meaning that the more times of adding noise, the greater the effect on model performance. Consequently, we need to handle the situation with multiple users when integrating DP into FL methods in real-life scenarios. To study the impact of the number of clients on the model performance with different methods and demonstrate the advantage of PPML-Omics, we tested the model performance by varying the number of clients on the same dataset. Since the integration of the DP mechanism requires adding noise to the gradient, increasing the number of clients had a significant impact on reducing the model performance of the FL-DP method (**Figure 2d**). As a consequence, a large number of clients in real life would be the biggest challenge for applying the FL-DP method [53]. However, the integration of the DR protocol could effectively weaken the performance degradation caused by increasing the number of clients, indicating that PPML-Omics could maintain better performance with a larger number of clients (≥ 10), making it a feasible solution in practical applications.

Another obvious advantage of PPML-Omics was the better data utility, meaning that PPML-Omics could retain a higher level of data utility while protecting data privacy. To show the data utility of PPML-Omics in the cancer classification task, in the ablation study, we compared the averaged accuracy and macro-F1 score on the testing dataset of the centrally trained method, the FL method, the FL-DP method, the FL-MHE method and PPML-Omics (**Figure 2c and Supplementary Table 2 and 5**). The FL method achieved slightly worse performance (accuracy=0.72830) than the centrally trained method (accuracy=0.73430) (*P* = 0.18577 for one-sided Student’s t-test). The FL-MHE method also achieved similarly good utility (accuracy=0.73902) as the centrally trained method (*P* = 0.76656 for one-sided Student’s t-test). Meanwhile, the FL-DP method under *ϵ*_*T*_ = 20 got the worst data utility (accuracy=0.62136). With the same end-to-end privacy guarantee (*ϵ*_*T*_ = 20), PPML-Omics showed significantly better utility (accuracy=0.67713) than the FL-DP method (accuracy=0.62136) (*P* = 0.0006 for one-sided Student’s t-test). In summary, PPML-Omics achieved good utility while preserving the privacy of gene expression.

Different cancer types have specifically expressed genes, which can be reconstructed by the attacker from the released models. Suppose that the attacker has auxiliary information, such as knowing that patient 1 is of cancer type A and has participated in the training of the released model. Thus, by performing the MIA, the attacker could roughly know the gene expression of patient 1, thus compromising the potential privacy of the patient as part of the model training data. To investigate the privacy protected by PPML-Omics and understand the biological meaning behind the JS divergence, we adopted the MIA for cancer classification models (**Methods**). In other words, we tried to optimize a gene expression vector that could give the highest prediction probability for a particular cancer type on the target model by using MIA. Then we analyzed the significantly expressed genes in the optimal gene expression vector. If the significantly expressed genes were cancer type-specific (with high expression in the target cancer type and low expression in other cancer types), then we could conclude that the target model leaked privacy. As shown in **Figure 2g**, the most significant genes reconstructed from the centrally trained method with H3R1D0 showed significantly different distribution (JS divergence = 0.13769, *P*_*KS*_ = 8.21642 × 10^−46^) on the z-score normalized real expression between two groups (solid lines for the target cancer type and dashed lines for other cancer types) compared to the one with PPML-Omics (*ϵ*_*T*_ = 20) (JS divergence = 0.00035, *P*_*KS*_ = 0.52341), suggesting that the MIA on the centrally trained method could reconstruct genes with significantly different expression levels (privacy leakage) on the target cancer type and other cancer types. In other words, based on the published centrally trained models, the attacker could reconstruct the corresponding specifically expressed genes for each cancer type. In contrast, it was impossible to accurately reconstruct the corresponding specifically expressed genes for each cancer type from the published models using PPML-Omics. A significant difference in the expression distribution of genes specifically reconstructed by the centrally trained method in the target cancer type and other cancer types could be observed (COADREAD: *P*_*KS*_ = 5.98454 × 10^−59^, KICH: *P*_*KS*_ = 3.59413 × 10^−34^ and UVM: *P*_*KS*_ = 2.14898 × 10^−131^) (**Figure 2h**), implying that genes reconstructed for each cancer type tended to have a higher expression in the corresponding cancer type. In contrast, the expression distribution of genes reconstructed by attacking PPML-Omics was very similar in the target cancer type and in other cancer types (COADREAD: *P*_*KS*_ = 0.03325, KICH: *P*_*KS*_ = 0.19957 and UVM: *P*_*KS*_ = 0.00144) (**Figure 2h**), implying that the genes reconstructed for each cancer type were not strongly correlated with the cancer type in terms of expression. A similar observation could be found across all cancer types that reconstructed genes by attacking the model trained with the centrally trained method showed significantly different expression levels between the target cancer type and other cancer types whereas reconstructed genes by attacking the model trained with PPML-Omics did not show a such significant difference (**Supplementary Section 2.3, Supplementary Figure 1, 2 and 3 and Supplementary Tables 3-4**). Overall, we showed that PPML-Omics protects genomic privacy by obfuscating sensitive gene identification.

Cryptography-based FL method could protect the security during the training phase, but this could also mean no protection at all, especially when the key leaks. We implemented an FL method based on multi-party homomorphic encryption (MHE), in which models and parameters were encrypted during the communication, training and inference phases using the homomorphic encryption (HE) method based on the Cheon-Kim-Kim-Song (CKKS) cryptographic scheme [61] (**Methods**). The clients and server have public and private keys for secure encryption and decryption. As shown in **Supplementary Tables 2 and 5**, the model trained with the FL-MHE method achieved similar accuracy (accuracy=0.73902) as the centrally trained method. Under the scenario of Application 1, the model could not be decrypted if assuming that the attacker could not obtain the public and private keys, thus the security and privacy are completely protected. Once the attacker obtained the public and private keys, the server model could be fully decrypted and the privacy leakage (JS divergence=0.10567, *P*_*KS*_ = 2.01988 × 10^−39^) was as serious as that of the centrally trained method as shown in **Figure 2c, Supplementary Figure 2 and Supplementary Table 5** even though the encryption and decryption processes introduced approximation errors and protected very little privacy. Meanwhile, the FL-MHE method required additional computational resources due to the encryption and decryption as shown in **Supplementary Table 2**. Overall, PPML-Omics showed a clear advantage over the FL-MHE method under our scenario in Application 1.

### Application 2. Clustering with scRNA-seq data

scRNA-seq is a revolutionary technology to quantify gene expression in thousands and even millions of cells in parallel, which is powerful to identify cell populations and dissect the composition of each population in biological samples [63]. Applying scRNA-seq on primary tumours is not only able to decipher tumour cell heterogeneity [64] but can also uncover the specificity of tumour micro-environment [65] in each patient, which may lead to a personalized treatment strategy [66]. For instance, immune infiltration in the tumour which could be determined by scRNA-seq, is a good response indicator to immune blockade-based therapy. Thus, information on cell populations is critical in protecting patients’ privacy when analyzing scRNA-seq data. Using the extracted features rather than the direct gene expression may avoid the leak of the specifically expressed gene while still harbouring the information for each cell population in the patients. Even so, obtaining accurate cell population results on patients’ tissue could also violate privacy. Here, we used the low-dimensional features extracted from gene expression vectors of scRNA-seq data by Auto-encoder as input for k-means clustering (**Figure 3a**). We then applied PPML-Omics to see how it protected patients’ privacy from the results of clustering, by evaluating the data utility and the privacy-preserving capability in terms of cell type classification and composition quantification based on clustering. The number and composition of cells included in the scRNA-seq data vary greatly depending on the sample. To assess the robustness of PPML-Omics for different sample sizes in the clustering task, we evaluated it on three benchmark datasets (Yan, Pollen, and Hrvatin) with varying cell numbers, ranging from 90 to 48,266 (**Supplementary Table 8**). Following Tran’s work [62], we compared the Adjusted Rand Index (ARI), Normalized Mutual Information (NMI), Cluster Accuracy (CA), and Jaccard index (JI) (**Methods**) on the three datasets with different methods (**Table 1**), including the centrally trained method, the FL method, the FL-DP method, the FL-MHE method and PPML-Omics, where a larger value means a better clustering result. For all three datasets, PPML-Omics achieved promising performance compared to the FL-DP method under all four evaluation metrics (Yan: *P* = 5.23 × 10^−3^, Pollen: *P* = 9.12 × 10^−4^, and Hrvatin: *P* = 1.01 × 10^−5^ from one-sided Student’s t-test) with the same privacy budget (*ϵ*_*T*_ =5). Also, the difference between the clustering result from PPML-Omics and that from the centrally trained model is relatively minor (Yan: *P* = 0.048, Pollen: *P* = 0.194, and Hrvatin: *P* = 0.012 from one-sided Student’s t-test), indicating that our framework even had the potential to approach the centrally trained method in clustering task. To ensure that the performance of PPML-Omics was at the same level as those commonly acknowledged tools in the clustering task with scRNA-seq data, we compared PPML-Omics with the existing state-of-the-art tools, including Seurat [67], SC3 [68], CIDR [69] and SINCERA [70] (**Supplementary Section 3.2 and 3.3, Supplementary Figure 4 and Supplementary Table 7**). PPML-Omics also achieved competitive utility, proving that our method achieved similar performance as the most commonly used tools for the clustering task. To further investigate the effect of FL, DP and DR protocol on the clustering task, we conducted an ablation study (**Figure 3b**)(centrally trained method at the first column, FL method (*ϵ*_*T*_ = 5, *ϵ*_*l*_ = 0.23) at the second column, FL-DP method (*ϵ*_*T*_ = 5, *ϵ*_*l*_ = 0.33) at the third column, and PPML-Omics at the fourth column) on these three datasets of different sizes and visualized the clustering results with uniform manifold approximation (UMAP) algorithm that projected the internal representations into a two-dimensional space. In conclusion, PPML-Omics could qualitatively and quantitatively compete or even outperform other methods in terms of utility.

**TABLE 1.**
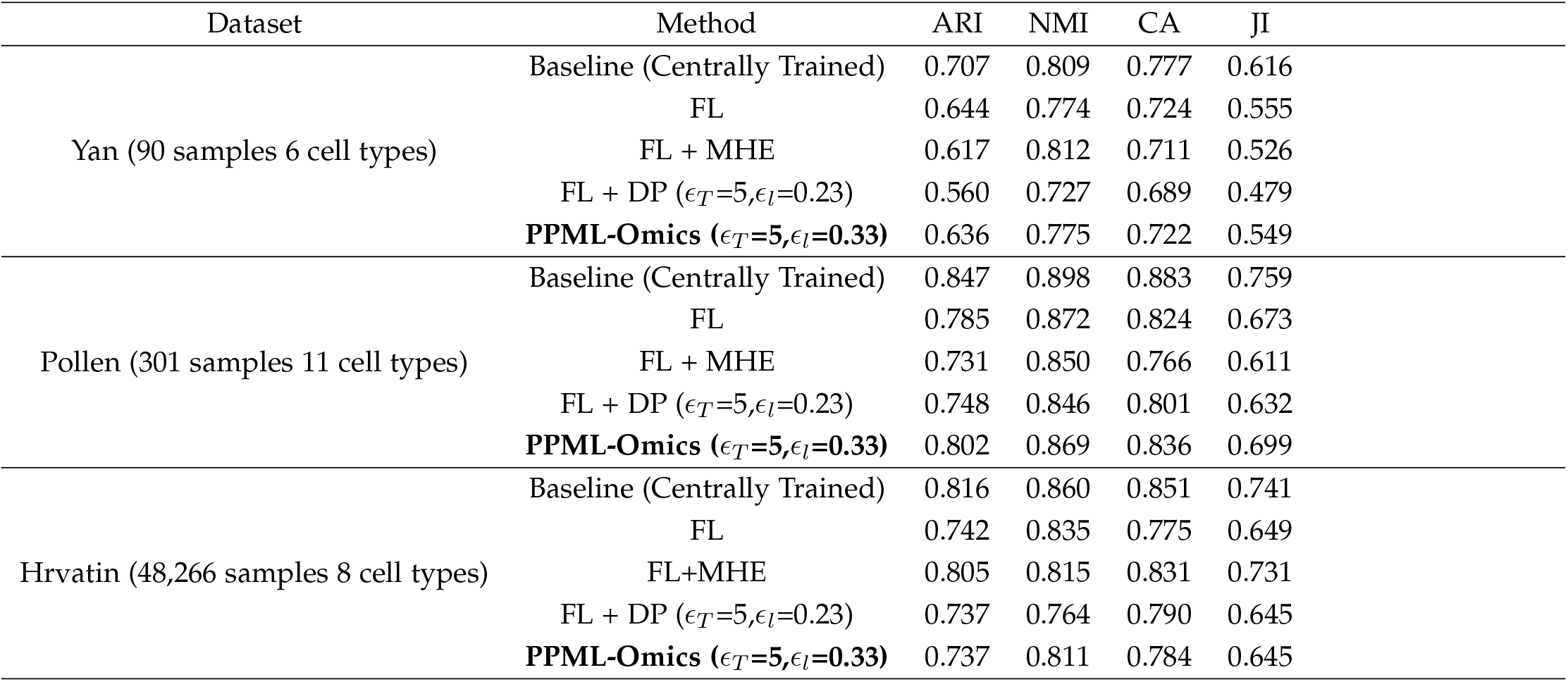
Comparison of clustering result on multiple scRNA-seq datasets [62]

**Fig. 3.**
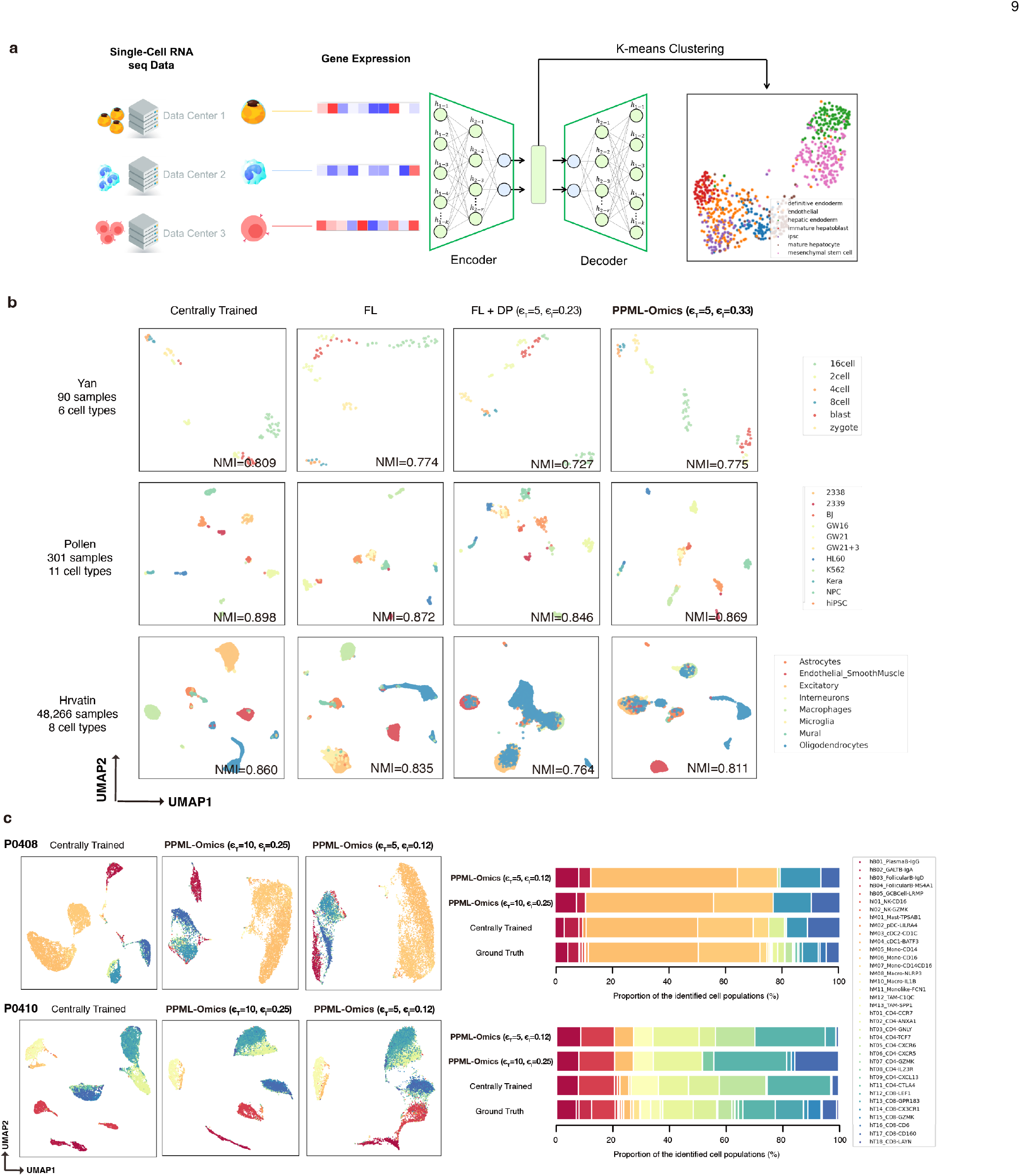
Results of clustering with scRNA-seq in Application 2. **a**, Architecture of the backbone (Auto-encoder). Gene expression vectors were fed into the Auto-encoder for feature reduction and selection, after which the low dimensional features were used for the K-means clustering. **b**, The clustering results visualized by the UMAP of three datasets with an increasing number of samples (Yan with 90 samples from 6 cell types, Pollen with 301 samples from 11 cell types, and Hrvatin with 48,266 samples from 8 cell types) generated by four methods, including the centrally trained method, the FL method, the FL-DP method and PPML-Omics. For these three datasets, all clustering results given by PPML-Omics showed a similar visual pattern to the centrally trained method, indicating the perfect utility of PPML-Omics. **c**, The clustering results visualized by the UMAP of two patients (P0408 and P0410) and the proportion plot of major clusters and sub-clusters generated by different methods, including the centrally trained method, and PPML-Omics with (*ϵ*_*T*_ = 10, *ϵ*_*l*_ = 0.25) and (*ϵ*_*T*_ = 5, *ϵ*_*l*_ = 0.12), indicating that PPML-Omics could protect patients’ privacy by removing the local information of sub-clusters.

When we obtain the scRNA-seq data of patients, we could learn the composition of the cells based on the clustering results. In extreme cases, suppose the clustering method is good enough and can 100% correctly distinguish different types of cells into clusters, it might potentially violate the patient’s privacy by leaking some sensitive sub-cell-types. Therefore, it is necessary to reasonably adjust the resolution of the clustering method to properly hide some small clusters (sub-cell-types) from the final clustering results to achieve privacy protection according to the prior privacy requirements. In other words, for a method, if we cannot observe some sensitive small clusters (sub-cell-types) in the final clustering results, then we could conclude that the method protects the patient’s privacy. Additionally, if the clustering results for major cell types are reasonable, we could conclude that the method preserves an acceptable degree of data utility while protecting privacy. To study the privacy-preserving power of PPML-Omics, we analyzed a public scRNA-seq dataset with 43,817 cells from 10 colon cancer patients [71], in which the cells were classified into 5 major cell types (hB: B cells, hI: Innate Iymphoid cells, hM: Myeloid cells, hT: CD4+ T cells and CD8+ T cells) and 38 sub-types **(Supplementary Section 3.5, Supplementary Tables 9-10**). We applied three methods for clustering, namely the centrally trained method and two of PPML-Omics with different noise levels (*ϵ*_*T*_ = 10, *ϵ*_*l*_ = 0.25) and (*ϵ*_*T*_ = 5, *ϵ*_*l*_ = 0.12). With two patients (P0408 and P0410) as examples, the major cell types were successfully identified with all three methods (**Figure 3c**), indicating that the overall clustering results of the three methods were reasonable, thus preserving the utility of the data. However, the clustering results from the three methods showed significant differences in the resolution of the sub-cell-types and we could use it as an evaluation of the privacy-preserving capability similar to the JS divergence in application 1. Comparing both our methods (*ϵ*_*T*_ = 10, *ϵ*_*l*_ = 0.25) and (*ϵ*_*T*_ = 5, *ϵ*_*l*_ = 0.12) with the centrally trained method, 4 sub-clusters (sub-cell-types) were undetected for P0408 and 6 sub-clusters (sub-cell-types) were undetected for P0410 **(Supplementary Tables 9-10**), suggesting that the local differences between sub-clusters (sub-cell-types) could be diluted due to the noise added in PPML-Omics and then those sub-cell-types could be integrated into major clusters, thus achieving privacy protection as a consequence.

Using a larger end-to-end *ϵ*_*T*_ means better utility and greater resolution on the clustering results (more sensitive sub-clusters could be observed), while using a smaller *ϵ*_*T*_ means that we could sacrifice a certain level of utility to protect sensitive sub-clusters in the clustering results. For potential users of PPML-Omics, the value of *ϵ*_*T*_ should be adjusted according to the needs of the actual application scenario to achieve both good utility and privacy protection as shown in **(Supplementary Figure 8**).

### Application 3. Integration of tumour morphology and gene expression with spatial transcriptomics

Sequencing data in Applications 1 and 2 could reveal much sensitive information about patients, and the potential risk of privacy disclosure is also very high for spatial transcriptomics as one of the highest-resolution sequencing technologies. In addition, spatial transcriptomics requires the usage of tissue images, which adds additional privacy leakage to some extent. ST-Net [56] is a machine learning model for predicting local gene expression from haematoxylin- and-eosin-stained pathology slides trained on a dataset of 30,612 spatially resolved gene expression data matched to histopathology images from 23 patients with breast cancer. Unlike the classical task of spatial transcriptomics, ST-Net predicts the spatially resolved transcriptome of tissue directly from tissue images, while the gene expression measured from spatial transcriptomics is used as the ground truth in the training phase. Here, to show that PPML-Omics realized competitive utility and to investigate how PPML-Omics protects privacy in those histopathology images on this task, we applied PPML-Omics to integrate tumour morphology and gene expression with spatial transcriptomics by incorporating the ST-Net model as the backbone network into PPML-Omics as shown in **Figure 4a**.

**Fig. 4.**
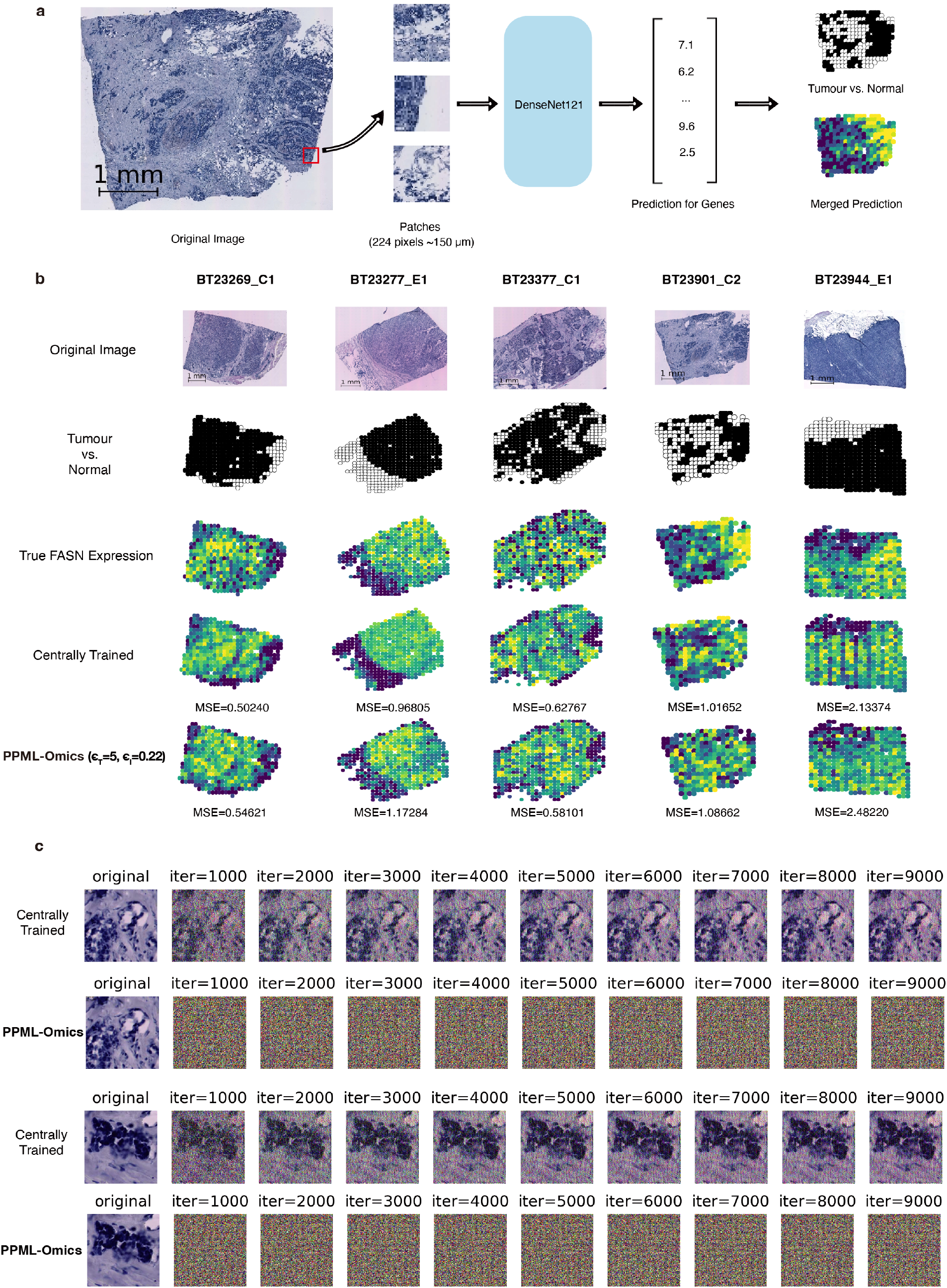
Results of integration of tumour morphology and gene expression with spatial transcriptomics in Application 3. **a**, Pipeline for predicting local gene expression from high-resolution tissue image referred by ST-Net. The patches (224×224) extracted from the original image were fed into the DenseNet121 and the local expression of selected cancer marker genes was predicted and compared with the real local expression (ground truth) from the spatial transcriptomics data. **b**, The results of five samples (BT23269 C1, BT23277 E1, BT23377 C1, BT23901 C2, BT23944 E1) showed the original image, the binary labels of the tumour (black) and normal regions (white), the real expression of *FASN* and predicted results by the centrally trained method and PPML-Omics (*ϵ*_*T*_ = 5, *ϵ*_*l*_ = 0.22). All predictions show a similar visual pattern with the ground truth and both the centrally trained method and PPML-Omics give similar good performance in terms of MSE. **c**, Image reconstruction attack with improved deep leakage from gradients (iDLG) on the centrally trained method and PPML-Omics (*ϵ*_*T*_ = 5, *ϵ*_*l*_ = 0.22).

To demonstrate that ST-Net under PPML-Omics has the same level of utility compared to pure ST-Net in predicting local gene expression from pathology slides, we visualized the prediction results and quantitatively compared the similarity between the predicted results and the ground truth by calculating the mean square error (MSE) as an evaluation metric, where a larger MSE means a worse prediction. From the visualization as shown in **Figure 4b**, both PPML-Omics (at the fifth row) and the centrally trained method (at the fourth row) obtained good results from a visual perspective compared to the ground truth (at the third row), effectively predicting the local expression of *FASN* and other cancer marker genes, including *HSP90AB1* and *PABPC1* (**Supplementary Section 4.2, Supplementary Figures 6-7**) with the histopathology images of 5 patients (BT23269 C1, BT23277 E1, BT23377 C1, BT23901 C2 and BT23944 E1) as inputs, which could be supported by the sequencing data from spatial transcriptomics (ground truth). Furthermore, the local gene expressions predicted by PPML-Omics and the centrally trained method were both aligned well to the tumour region (black) and normal region (white) annotated in the annotation (at the second row), indicating that the ST-Net integrated with PPML-Omics still had the power to predict the local gene expression accurately from tissue images. In addition, with the same small privacy budget *ϵ*_*T*_ = 1 and 5, the prediction of PPML-Omics was significantly more accurate (MSE=0.98808 at *ϵ*_*T*_ = 1 and MSE=0.93005 at *ϵ*_*T*_ = 5) than the FL-DP method (MSE=1557.54465 at *ϵ*_*T*_ = 1 and MSE=1.29931 at *ϵ*_*T*_ = 5) with *P* = 8.06039 × 10^−29^ and *P* = 3.10063 × 10^−14^ for one-sided Student’s t-tests respectively and achieved close performance of the centrally trained method (MSE=0.90904) (**Table 2 and Supplementary Table 13**). Thus, PPML-Omics achieved good utility on Application 3.

**TABLE 2.** Comparison of different methods in the integration task of spatial gene expression and tumour morphology with spatial transcriptomics.

| Method | Averaged Testing MSE |
| --- | --- |
| Baseline (ST-Net, Centrally Trained) | 0.90904 |
| Baseline + FL | 0.93120 |
| Baseline + FL + MHE | 0.90904 |
| Baseline + FL + DP ($\epsilon_T=0.1, \epsilon_l=0.003$) | NA |
| Baseline + FL + DP ($\epsilon_T=0.5, \epsilon_l=0.016$) | NA |
| Baseline + FL + DP ($\epsilon_T=1, \epsilon_l=0.032$) | 1557.54465 |
| Baseline + FL + DP ($\epsilon_T=5, \epsilon_l=0.160$) | 1.29931 |
| **PPML-Omics ($\epsilon_T=0.1, \epsilon_l=0.004$)** | 3.22789 |
| **PPML-Omics ($\epsilon_T=0.5, \epsilon_l=0.022$)** | 1.08788 |
| **PPML-Omics ($\epsilon_T=1, \epsilon_l=0.045$)** | 0.98808 |
| **PPML-Omics ($\epsilon_T=5, \epsilon_l=0.225$)** | 0.93005 |

It is well known that medical imaging data contains much potentially sensitive information [11], such as tissue patterns and lesions, which could compromise patients’ privacy. Several studies have shown that machine learning models trained based on medical images remember much information about the images in the training dataset, and attackers could perform image reconstruction attacks with the published machine learning models to obtain the original training images [72], [73], [74]. To see whether PPML-Omics protects privacy in the histopathology images compared to the centrally trained method, we used the iDLG [75] (**Methods**) to simulate an attacker reconstructing the training images by stealing the gradients passed between machines under the FL framework (approximating a centrally trained method when the number of clients is 1). In each attack, we initialized a noisy input (dummy data), used the local gene expression as the ground truth, computed the prediction of the model on the noisy input and calculated the gradient. Then, we updated the noisy input with the gradient. In other words, same as in Application 1, we were trying to optimize an input image that could get the most similar prediction results to the ground truth. We performed the iDLG attack separately for the centrally trained method and PPML-Omics and showed the input image every 1000 iterations, as shown in **Figure 4c**. Strikingly, in terms of visual results, the iDLG attack could effectively reconstruct the training images with the centrally trained method, while in PPML-Omics, the iDLG attack was effectively blocked due to the addition of noise to the gradients with the DP mechanism. Overall, PPML-Omics protected privacy by blocking the reconstruction of sensitive histopathology images.

### Theoretical proof of the privacy-preserving power of PPML-Omics

With the three applications above, we have empirically demonstrated our method’s privacy-preserving capability and utility. We further theoretically proved the privacy-preserving capability of PPML-Omics with the DP notation. The central DP (CDP) model and the local DP (LDP) model are two commonly acknowledged models with the notation of DP. In the CDP model, a server trusted by users collects users’ raw data (e.g. local updates) and executes a private mechanism for deferentially private outputs. The privacy-preserving goal is to achieve indistinguishability for any outputs w.r.t. two neighbouring datasets that differ by replacing one user’s data. The definition of differential privacy (**Definition 4 in Supplementary Section 1.2**) requires that the contribution of an individual to a dataset has not much effect on what the adversary sees. Compared to CDP, LDP is a stronger notion of privacy. In LDP, each user’s data is required to be perturbed to protect privacy before collection. The CDP model assumes the availability of a trusted analyzer to collect raw data, while the LDP model does not rely on any trusted party because users send randomized data to the server. The CDP model protects the calculation result in the analyzer, so users need to trust the central server and send raw data to the server, which allows greater accuracy but requires a trusted analyzer which is impractical in most real cases. In the LDP model, we protect the data information in the single local device, so users only need to trust their single device and randomize local data before sending them to the analyzer. Though a trusted central analyzer is not required, the utility of the method is limited because we do lots of randomness on local data. Therefore, the advantage of using the DR protocol (**Definition 6 in Supplementary Section 1.4**) is that we could balance the strength in both CDP and LDP, i.e., good performance of accuracy in the CDP model and strong privacy in the LDP model without relying on any trusted central party.

Under the FL, each client trains its model locally and sends the update in the form of gradients to a central server that would aggregate those updates into the central model. After updating the central model, the central server broadcasts the new model weight to all clients for updating all clients (**Algorithm 1**). With the integration of DP in the FL framework, we manually chose several end-to-end *ϵ*_*T*_ and set *δ* as discussed in **Methods**. Given *ϵ*_*T*_ and *δ*_*T*_, we applied the analytic Gaussian mechanism to calculate the optimal *σ* for perturbation [76] as in **Algorithm 2**. In other words, we could calculate the amount of noise that needs to be added based on the given *ϵ*_*T*_ and *δ*_*l*_. The privacy guarantee of the analytic Gaussian mechanism was given in (**Theorem 1 in Supplementary Section 1.4**). Thus, we applied the analytic Gaussian mechanism on a single client with the privacy guarantee proved to realize the DP mechanism.

Based on the post-processing property (**Lemma 1 in Supplementary Section 1.4**), the protocol 𝒫 (defined in **Definition 6 in Supplementary Section 1.4**) achieves the same privacy level as ℳ (defined in **Lemma 1 in Supplementary Section 1.4**) because 𝒜 is executed by an untrusted analyzer, without protecting users’ privacy. We wanted to obtain ℳ = 𝒮 ° ℛ^*n*^ that could satisfy (*ϵ*_*c*_, *δ*_*c*_)-DP, meaning that we achieved the same privacy guarantee as the CDP. Thus, we focused on analyzing the indistinguishability for ℳ (*X*) and ℳ (*X*^′^) where *X* and *X*^′^ differ in one client’s local vector such that we achieved privacy-preserving on the client-level. [77] proved that the privacy of ℳ could be amplified. In other words, when each user applies the local privacy budget *ϵ*_*l*_ in ℛ, ℳ can achieve stronger privacy of (*ϵ*_*c*_, *δ*_*c*_)-DP with *ϵ*_*c*_ *< ϵ*_*l*_. Hence, the DR protocol has a larger privacy budget for a local single client and needs less noise to achieve the same privacy model compared with the LDP model. In practice, we defined a privacy budget *ϵ*_*c*_ in a single epoch, then we amplified this budget to the local client model with the local privacy budget *ϵ*_*l*_.

Given target privacy parameters 0 *< ϵ*_*T*_ *<* 1 and 0 *< δ <* 1, to ensure (*ϵ*_*T*_, *δ*_*T*_)-DP over *T* mechanisms, it suffices that each mechanism is (*ϵ*_*c*_, *δ*_*c*_)-DP, where 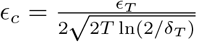 and 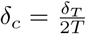. From advanced composition theorem and the Corollary (**Supplementary Section 1.4**), we can guarantee that Algorithm 2 satisfies (*ϵ*_*T*_, *δ*_*T*_)-DP after T epochs.

## Discussion

Overall, PPML-Omics is universal, meaning that PPML-Omics could be integrated with any machine learning model and applied to various biological problems. Thus, we applied PPML-Omics to three different machine learning models from simple to complex, including FCN, Anto-encoder, and DenseNet-121. Applying PPML-Omics to more complex deep learning models means that more computational resources are required to add noise, which may introduce more uncertainty in the performance of the models. Hence, whether PPML-Omics can be applied to more complex machine learning models and used to protect privacy in more complex data requires further research in the future.

The integration of the DP mechanism requires additional computational resources, such as more GPU memory consumption and computation time, for calculating the magnitude of the noise and the gradient calculation with noise. However, we demonstrated in our simulated experiments that the computational burden imposed by PPML-Omics was insignificant compared to centrally trained methods, implying that potential users could easily deploy PPML-Omics for privacy-preserving omic data analysis without upgrading their existing computational devices.

It is commonly acknowledged that privacy protection and utility are two conflicting objectives. Thus it is challenging to find a balance that allows the model to protect privacy without overly damaging the usability of the data. Therefore, PPML-Omics balances the privacy-preserving capabilities with utility as much as possible, while also leaving some of the decision-making to our potential users. Depending on the user’s actual needs, the user can adjust the end-to-end privacy budget (*ϵ*_*T*_) in the method during the training phase or select the released model trained with PPML-Omics under different *ϵ*_*T*_ to achieve different levels of privacy protection. We acknowledge the fact that PPML-Omics could not automatically select *ϵ*_*T*_ for potential users for specific biological problems but requires the user to manually select *ϵ*_*T*_ is a major weakness of PPML-Omics. But there are potential reasons for this as different biological data have different characteristics as well as various inherent noise, and different deep learning models could tolerate different levels of noise. For example, for the three applications, we selected different values of *ϵ*_*T*_ in order to achieve appropriate privacy protection (e.g., *ϵ*_*T*_ = 20 in Application 1, *ϵ*_*T*_ = 5 in Application 2, and *ϵ*_*T*_ = 5 in Application 3). It may take more time for users to try different values of *ϵ*_*T*_ to choose an appropriate one that fits the privacy requirement the most when applying PPML-Omics in practice. We also provided the relationship between various evaluation metrics and the value of *ϵ*_*T*_ in all three applications as a reference (**Figure 2e,f, Supplementary Figure 8 and 9**).

In summary, we have proposed a secure and privacy-preserving machine learning method by designing a decentralized version of the differential private federated learning algorithm (PPML-Omics). Besides, we have applied this method to analyze data from three representative omic data analysis tasks, which are solved with three different deep learning models, revisited and addressed the privacy concern in the cancer classification from TCGA with bulk RNA-seq, clustering with scRNA-seq, and the integration of spatial gene expression and tumour morphology with spatial transcriptomics. Moreover, we examined in depth the privacy breaches that existed in all three tasks through privacy attack experiments and demonstrated that patients’ privacy could be protected by PPML-Omics. Additionally, we proved the privacy-preserving capability of PPML-Omics theoretically, suggesting the first mathematically guaranteed method with robust and generalizable empirical performance in protecting patients’ privacy in omic data. We believe that PPML-Omics will attract the attention of future researchers on biological issues regarding data privacy, and also try to protect data privacy by applying PPML-Omics in research. Our method’s modularized and extendable nature has great potential to be developed collaboratively for different biological tasks and deep learning models, and will shed light on the application and development of deep learning techniques in the privacy protection of biological data.

## Methods

### Dataset preparation

For Application 1, we used the Cancer Genome Atlas (TCGA) dataset, accessible through the Genomic Data Commons Data Portal and built our methods across 37 cancers with a total number of 13,057 patients. Each patient had the expression vector of 20,531 genes, and each expression vector was re-scaled in logarithm with 10 according to (1), to speed up learning and lead to faster convergence. Also, the normalization procedure ensured that the gene expression contributing to the model was on an equal scale. Patients were randomly divided into a training dataset (80%) and test dataset (20%) and the random split was repeated 5 times (**Supplementary Section 2.1**).

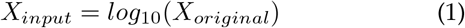

For Application 2, to assess the representation from our privacy preserved model on the scRNA-seq data, we tested our model on 3 published datasets with different sample sizes, all of which have expert-annotated labels from scDHA [62]. The Yan [78] dataset refers to the human preimplantation embryos and embryonic stem cells. In this dataset, 90 cells were sequenced with the Tang protocol. We first log-transformed the gene expression values selected for the highly variable genes. Finally, we scaled the dataset to unit variance and zero mean. The Pollen [79] dataset was sequenced with the SMARTer protocol. It contains 301 cells in the developing cerebral cortex from 11 populations. We downloaded it from the Hemberg Group’s website (https://hemberg-lab.github.io/scRNA.seq.datasets/human/tissues/pollen) and removed the low-quality data and cells with more mitochondrial genes and spike-in RNA. Then we selected highly variable genes after logarithmic transformation. The Hrvatin [80] dataset contains 48,266 cells from 6-8 week old mice, which were sequenced by DropSeq. After filtering the low-quality data, we performed the aforementioned logarithmic transformation, normalization per cell count, and highly variable gene selection steps. To further validate the privacy-preserving ability of PPML-Omics, we selected the scRNA-seq data on immune and stromal populations from colorectal cancer patients [71]. We visualized the global and local clusters regarding their specific macrophage and conventional dendritic cell (cDC) subsets. The single-cell data were selected from patients with informed mechanisms of Myeloid-Targeted therapies in colon cancer, which can be found in Data Availability. After quality control, each patient has around 5,000 cells and corresponding 13,538 genes (**Supplementary Section 3.1**).

For Application 3, to test the capability of our method in spatial transcriptomics, we used a spatial transcriptomics dataset of 30,612 spots, including 23 patients with breast cancer with four subtypes: Luminal A, Luminal B, Triple-negative, and HER2-positive [56]. There were 3 high-resolution images of H&E staining tissue and their corresponding gene expression data for each patient. The number of spots for each replicate ranged from 256 to 712 depending on the tissue size and the diameter of a single spot was 100 *μm* arranged in a grid with 200 *μm* as the centre-to-centre distance. There were 26,949 mRNA species detected across the dataset and each spot was represented as a 26,949-dimensional vector containing the elements denoted by the number of gene counts (**Supplementary Section 4.1**).

### Deep learning model design and training

All codes were implemented using the PyTorch library [81] and processed on one machine with 2 NVIDIA V100 GPUs and 252G RAM. All experiments were repeated on the pre-split training and testing batches until fully converged on the testing dataset, and all reported performance was averaged over five random splits.

For Application 1, we applied a fully connected neural network with 3 hidden layers as our benchmark machine learning network for PPML-Omics. The gene expression vector was fed into the network, and a ReLU layer was applied after each hidden layer to provide non-linearity. Finally, a Soft-max layer was attached to the last hidden layer to get the final prediction. With PPML-Omics, the network was trained in a federated, secured and privacy-preserving procedure to guarantee the utility while preserving privacy at the same time. The multi-class cross-entropy loss defined the loss function:

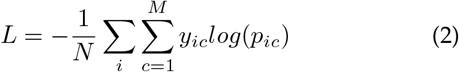

where *M* represents the number of cancers (classes), which is 37 for this task. Correspondingly, *y*_*ic*_ ∈ {0, 1} represents the classification result for the *i*-th sample, with 1 for a positive prediction, and 0 for a negative prediction. *p*_*ic*_ stands for the probability of the *i*-th sample to be in class *c*.

For Application 2, the unsupervised framework for scRNA-seq clustering is composed of two steps. We first trained the Auto-encoder with the fixed number of gene expression value *y* as the input and outputed *ŷ* with the same size of the input. We denoted *d* as the dimension of each patient’s gene expression vector and *N* as the number of clients. We then optimized the Auto-encoder with the Mean squared error (MSE) loss:

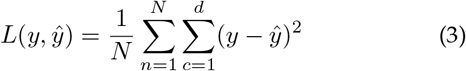

Then we extracted the central feature vector and clustered with *k*-means clustering

For Application 3, we used ST-Net [56] as our baseline deep learning network and integrated it into PPML-Omics. In the ST-Net, we used DenseNet-121 to detect the fine-grained spatial heterogeneity within the tumour tissue. Small patches (224×224 pixels) were extracted from the whole-slide images (∼10000×10000 pixels) centred on the spatial transcriptomics spots, and each patch went through the pre-trained DenseNet-121 network for training and predicting the pre-selected 250 genes with the highest mean expression.

### Performance assessment

To assess the performance of models in Application 1, we adopted metrics including accuracy and macro-F1, which are defined as below:

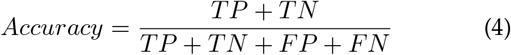

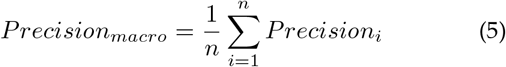

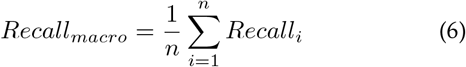

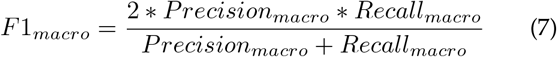

where TP, TN, FP and FN stand for true positive, true negative, false positive and false negative, *y*_*i*_ is the predicted value, 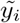 is the real value, and *n* is the total number of samples.

To evaluate PPML-Omics on Application 2, we applied clustering Adjusted Rand Index (ARI) score, Normalized Mutual Information (NMI), Cluster Accuracy (CA), and Jaccard index (JI) to evaluate our method’s utility comprehensively. The evaluation metrics between the predicted cluster *B* with the ground truth cluster *A* in ARI, NMI, CA, and JI are defined below:

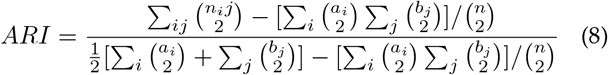

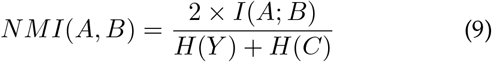

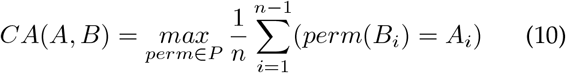

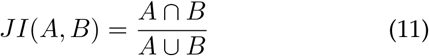

where A is the class labels, B is the cluster labels, *H*(·) is the Entropy and *I*(*A*; *B*) is the Mutual Information between A and B. In the definition of Cluster Accuracy (CA): *P* stands for the set of all permutations in [1:K] where K is the number of clusters, and *n* is the sample size.

To measure the similarity between the predicted gene expression and the ground truth in Application 3, we used the mean squared error (MSE) as the evaluation metric:

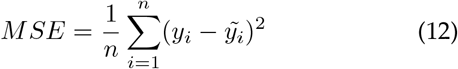

### Implementation of federated learning framework

As shown in **Algorithm 1**, the federated learning framework aimed at ensuring data privacy and security along with the improvement of the AI model based on joint data sources from multiple clients around the world. We first initiated multiple data sources as our clients. Since the essence of federated learning is the union of samples, each client first needs to download the model from the server and initiate their client model with the server weight. Then each participant can use local data to train the model, calculate their gradient, and upload it to the server. The server needs to aggregate the gradient of each client to update the server model parameters. In this process, each client is treated with the same and complete model, and there is no communication and no dependence among clients. Therefore, each client can also make independent predictions during the prediction.

### Implementation of differential private model training

To add noise to the gradient, we used the calibrate analytic Gaussian mechanism to calculate the value of *σ* in the Gaussian distribution based on the values of *ϵ* and *δ*. In each epoch, after computing the gradient update for each client, we clipped the gradient to ensure a finite upper and lower bound and then added noise to each value in the gradient that fitted the previously defined Gaussian distribution as shown in **Algorithm 2**. A common paradigm for approximating a deterministic real-valued function *f* : *D* → *R* with Gaussian differentially private mechanism is via additive noise calibrated to *f*’s sensitivity *S*_*f*_, which is defined as the maximum of the *l*_2_-norm ‖*f* (*D*) − *f* (*D*^′^)‖_2_ where *D* and *D*^′^ are neighbouring inputs. Proving the differential privacy guarantee in the SGD algorithm requires bounding the influence of each sample on the gradient. Since there was no prior bound on the size of the gradients, we clipped each gradient in *l*_2_ norm by *C*.

#### Algorithm 1. Federated learning

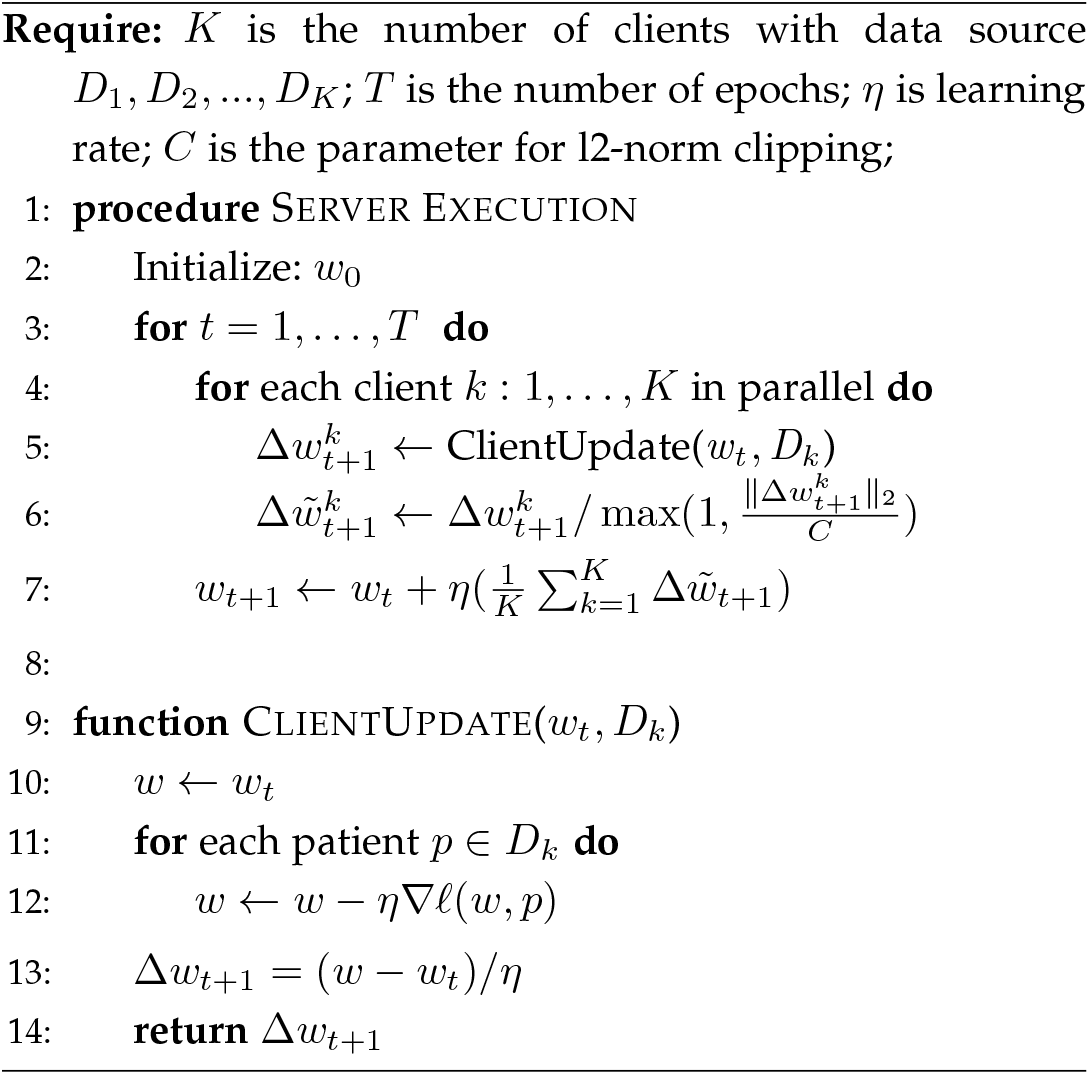

### Implementation of DR protocol

In a practical scenario, before the end of each epoch, each client sends the gradient information to the server, while the server knows where each gradient information comes from, providing the possibility of privacy leakage. With DR protocol, we can still ensure that the gradients received by the server are randomly shuffled. As shown in **Algorithm 2**, in each epoch of training, after the DR mechanism, all clients may not hold their own original gradients, but rather gradients from a randomly paired client. Then all clients upload their gradients to the server.

### *σ* selection with analytic Gaussian DP mechanism

In order to achieve end-to-end (*ϵ*_*T*_, *δ*_*T*_)-DP for T iterations in total with PPML-Omics, we need to achieve (*ϵ*_*c*_, *δ*_*c*_)-DP for the server in each epoch where 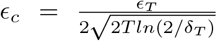 and 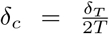, and further achieve (*ϵ*_*l*_, *δ*_*l*_)-DP for each client at each epoch, where 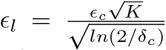 and 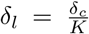. With the values of *ϵ*_*l*_ and *δ*_*l*_, we could calculate the standard deviation *σ* of Gaussian noise needed in each epoch to each gradient transmitted from each client by using the analytic Gaussian mechanism of [76], where *σ* = *CalibrateAnalyticGaussianMechanism*(*ϵ*_*l*_, *δ*_*l*_). FL-DP method, we could calculate 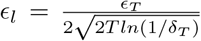 and 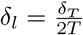.

#### Algorithm 2. PPML-Omics

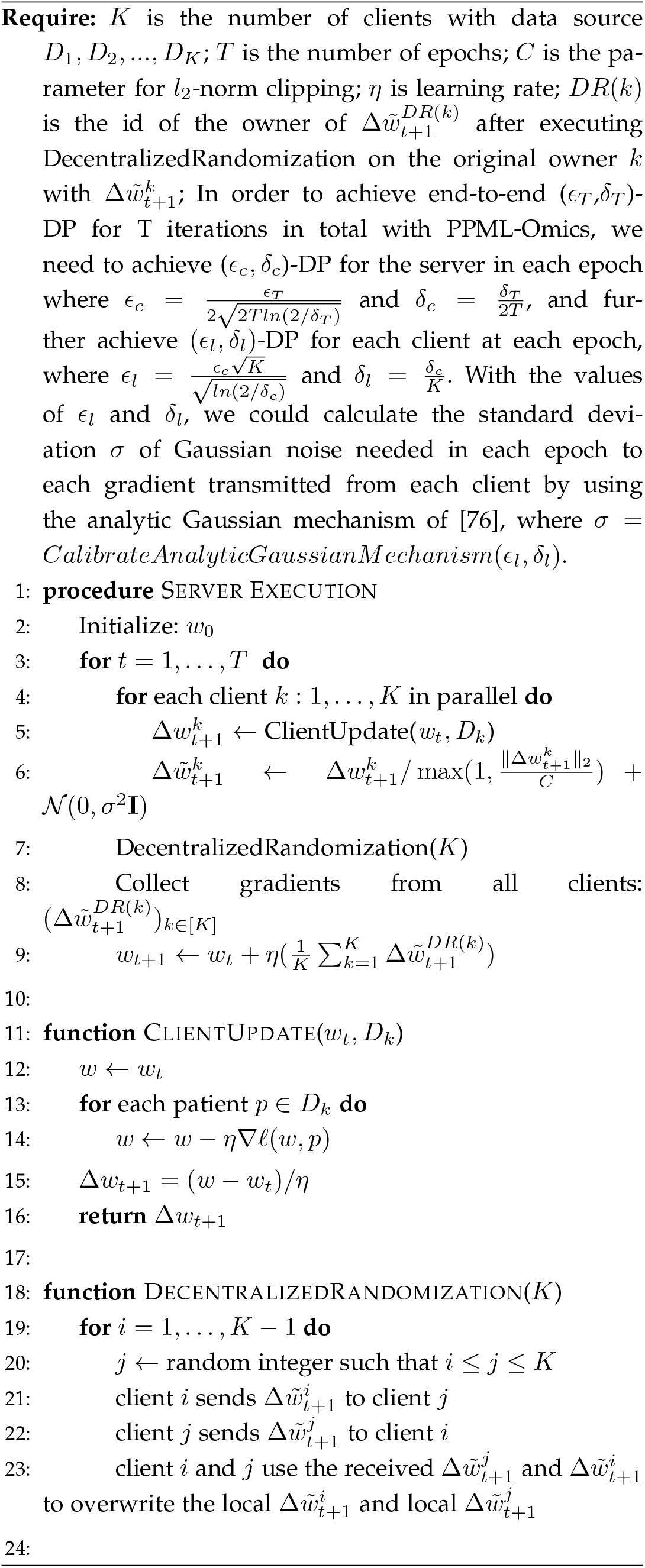

Let *f* : 𝒳^*n*^ → ℝ ^*k*^ be a function with global *ℓ*_2_-norm sensitivity Δ. Suppose *ϵ >* 0 and 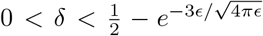. If the Gaussian mechanism *A*(*D*) = *f* (*D*) + *Z* with *Z* ∼ 𝒩(0, *σ*^2^𝕀_*k*_) is (*ϵ, δ*)-DP then 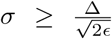. When 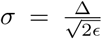 in analytic Gaussian DP mechanism, from Theorem 1 in supplement, the mechanism will be (*ϵ, δ*)-DP with 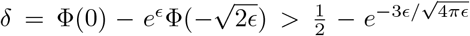.Thus, it is impossible to achieve (*ϵ, δ*)-DP with 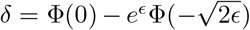 where without increasing the variance of perturbation.

We could use the above theorem to get the minimal noise to add for the given *ϵ*_*l*_ for each client at each epoch.

### Hyperparameter selection in DP mechanism

Regarding the parameters in the DP mechanism, we selected the proper value **C** for *l*_2_-norm clipping based on statistically observing the distribution of all elements in gradients during the training phase for the centrally trained model, thus for estimating the sensitivity of the training procedure of PPML-Omics. The selection of the privacy budget *ϵ* was a tricky task, as commonly adopted, such as in the handbook ‘Disclosure Avoidance for the 2020 Census: An Introduction’ [59] and Jayaraman et al. [82], we tested *ϵ* from 0.1, 0.5, 1, 5, 10, 20, 30, 40, 50 and measured the utility against different end-to-end *ϵ*_*T*_ in for all applications as a reference the potential users of PPML-Omics. To choose the proper value of *δ*, we performed the grid search from 0 to 1/*K*, where *K* is the number of clients.

### Hyperparameter and model selection in deep learning

To ensure that all models had a fair chance of learning a useful representation in all tasks, we trained multiple instances of each model (FCN for Application 1, Auto-encoder for Application 2 and ST-Net for Application 3) using a grid search of hyperparameter settings, including learning rate, epochs, number of hidden layers, number of filters, filter size, batch size, momentum, initial weight, weight decay and keep probability. Then, we selected model instances based on their training performance.

### Model inversion attack for cancer classification models

ML model was abused to learn sensitive genomic information about individuals, which was shown in a case study of linear classifiers in personalized medicine by Fredrikson et al. [83]. Thus, we implemented the model inversion attack (MIA) proposed by Fredrikson et al. [84] to extract sensitive information from the trained models with our PPML-Omics method under different privacy budgets to address the privacy concerns. Regarding our tasks, we retreated the MIA as an optimization problem: to find the input that maximizes the returned class confidence score [84], which was achieved by using gradient descent along with modifications specific to this domain as shown in Algorithm 3 (**Supplementary Section 2.3**).

We reconstructed the original gene expression with the MIA for each cancer for all pre-trained models. For each cancer, we applied MIA to reconstruct the gene expression vector for the targeting cancer and selected genes with the highest reconstructed expression levels (≥0.8) as the most significant reconstructed gene features specific to each cancer. Then we split all samples of 37 cancers into 2 groups (one only included the samples of the targeting cancer type and the other one included all remaining samples from other cancer types) and plotted the distribution of the z-score normalized expression of the previously selected genes of the real expression data in both groups (solid lines for samples of the target cancer type and dashed lines for other cancer types).

#### Algorithm 3. Model Inversion Attack (MIA)

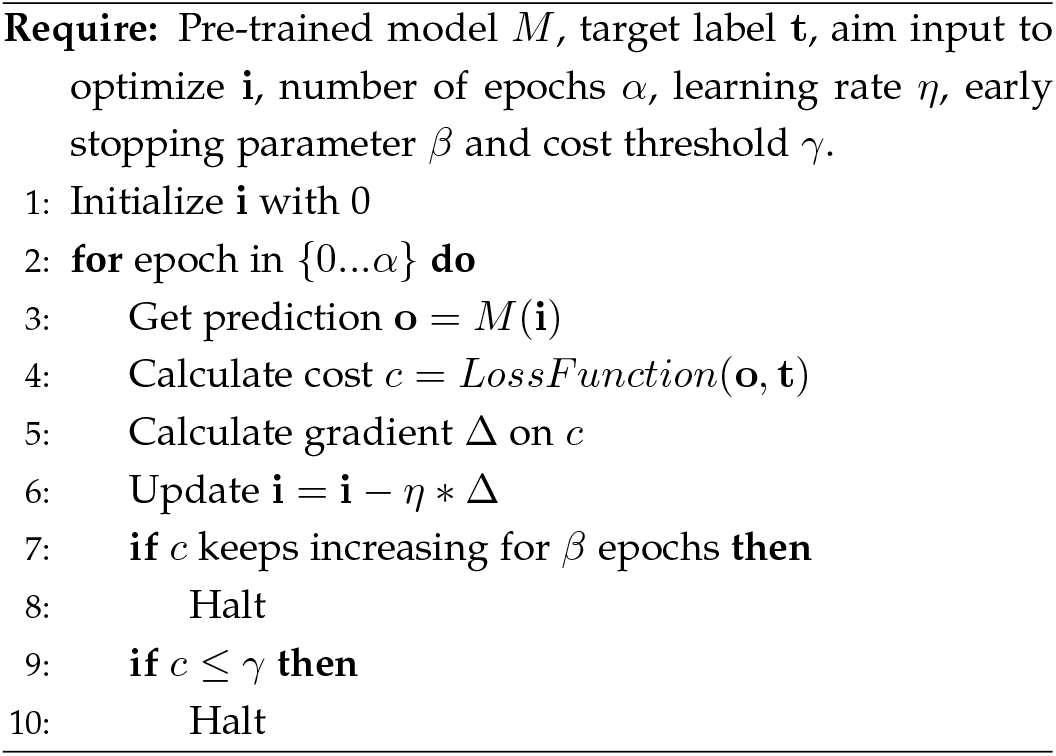

### iDLG reconstruction attack

Sharing gradients during the training of machine learning networks could potentially leak private data. Zhao et al. presented an approach named improved deep leakage from gradient (iDLG) as shown in **Algorithm 4**, which showed the possibility to obtain private training data from the publicly shared gradients. In integrating tumour morphology and gene expression with spatial transcriptomics, we synthesized the dummy data and corresponding labels with the supervision of shared gradients and optimized the dummy data with iDLG to obtain the one similar to the private training data.

#### Algorithm 4. iDLG

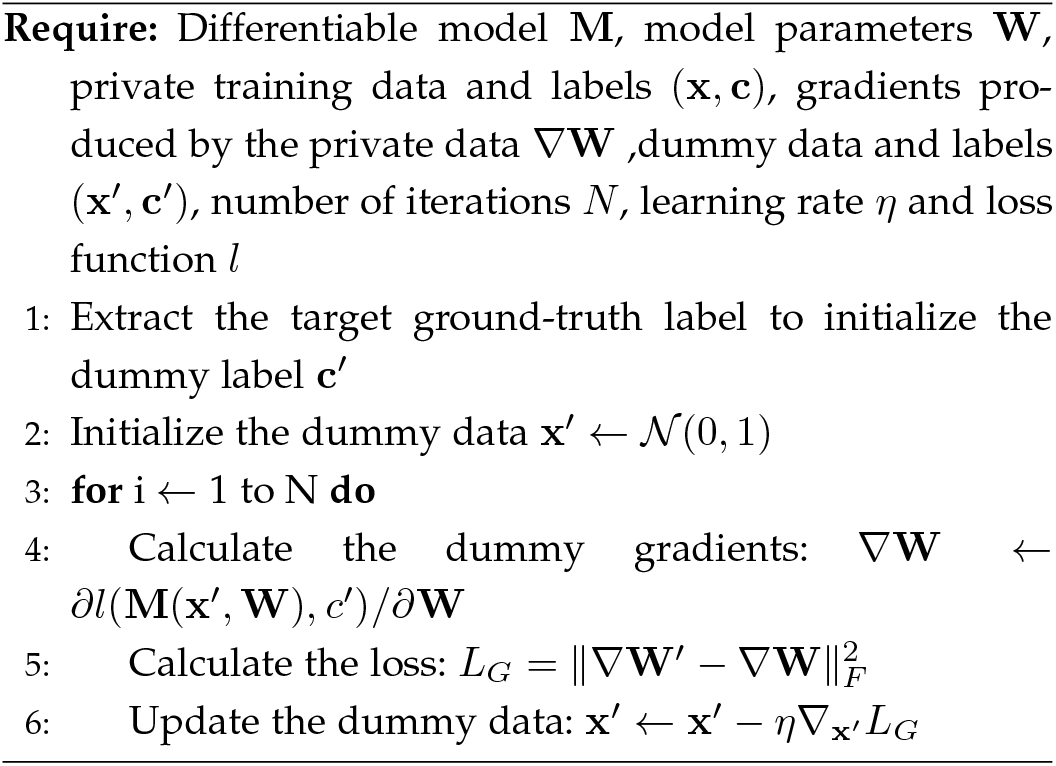

### Quantification of privacy leakage

For Application 1, to investigate the significance of the results of gene reconstruction attacks, we used Kullback–Leibler (KL) divergence, Jensen-Shannon (JS) divergence and Kolmogorov–Smirnov (KS) test as shown below to characterize the degree of privacy leakage by calculating the divergence between two distributions of the expression level of the reconstructed genes in the target cancer type and other cancer types. Larger JS divergence values and smaller p-values of the KS test indicated more significant differences between the expression levels of the reconstructed signature genes in the target cancer type and other cancer types, suggesting a severe privacy leakage.

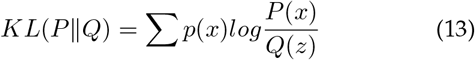

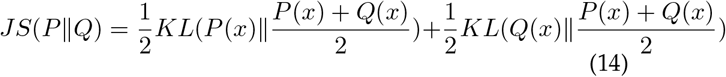

where P(x) and Q(x) are two probability distributions.

For Application 2, given a method, if we cannot observe some targeted small clusters (sub-types) in the final clustering and the proportion results, then we could conclude that the method protects the patient’s privacy. Additionally, if the clustering results for major cell types are correct, we could conclude that the method preserves a reasonable degree of usability while protecting privacy.

PPML-Omics avoids reconstruction attacks on medical images. Thus we evaluated the privacy-preserving capability by visually comparing the reconstructed image and the raw image after the iDLG attack on different methods in Application 3.

### Multi-party homomorphic encryption (MHE)

To compare PPML-Omics (DP-based solution) with the solution in the cryptographic track, we implemented an FL method with multi-party homomorphic encryption (MHE) based on the Cheon-Kim-Kim-Song (CKKS) cryptographic scheme [61] that provides approximate arithmetic over vectors of complex numbers in the training phase of FL method with TenSEAL [85], which is a library for doing homomorphic encryption operations on tensors, built on top of Microsoft SEAL.

## Author contributions

J.Z. and S.C. contributed the original idea, model design, experimental analysis and wrote the manuscript. Y.W contributed the theoretical proof and wrote the manuscript. The review and editing of the manuscript and some supplementary visualizations were carried out by H.L., B.Z., L.X., Y.H., Z.X., Z.L., N.C. and W.H. The entire project was supervised by X.G. and D.W..

## Data and materials availability

Data for the cancer classification with bulk RNA-seq is based upon the one generated by the TCGA Research Network: https://www.cancer.gov/tcga. The datasets for scRNA-seq clustering analysis are available at GEO accession ID: Yan (GSE36552), Pollen (SRP041736), and Hrvatin (GSE102827). The rds files are also available on the SCDHA software availability page (https://bioinformatics.cse.unr.edu/software/scDHA/resource/Reproducibility/Data/). scRNA-seq data of patients are available at GEO accession ID: GSE146771. For the spatial transcriptomics dataset, there are 30,612 spots, including 23 patients with breast cancer with four subtypes: Luminal A, Luminal B, Triple-negative and HER2-positive. There were 3 high-resolution images of H&E staining tissue and their corresponding gene expression data for each patient. All images and processed data are available at http://www.spatialtranscriptomicsresearch.org.

## Code availability

Code for this paper is available at https://github.com/JoshuaChou2018/PPML-Omics.

## REFERENCES

[1] Y. Joly, S. O. M. Dyke, B. M. Knoppers, and T. Pastinen, “Are Data Sharing and Privacy Protection Mutually Exclusive?” Cell, vol. 167, pp. 1150–1154, 11 2016.

[2] K. Tomczak, P. Czerwińska, and M. Wiznerowicz, “The cancer genome atlas (tcga): an immeasurable source of knowledge,” Contemporary oncology, vol. 19, no. 1A, p. A68, 2015.

[3] M. Caulfield, J. Davies, M. Dennys, L. Elbahy, T. Fowler, S. Hill, and K. Woods, “The 100,000 genomes project protocol,” Genomics Engl. P, vol. 3, pp. 1–112, 2017.

[4] H. A. Lewin, G. E. Robinson, W. J. Kress, W. J. Baker, J. Coddington, K. A. Crandall, R. Durbin, S. V. Edwards, F. Forest, M. T. P. Gilbert et al., “Earth biogenome project: Sequencing life for the future of life,” Proceedings of the National Academy of Sciences, vol. 115, no. 17, pp. 4325–4333, 2018.

[5] M. Alser, H. Hassan, H. Xin, O. Ergin, O. Mutlu, and C. Alkan, “Gatekeeper: a new hardware architecture for accelerating prealignment in dna short read mapping,” Bioinformatics, vol. 33, no. 21, pp. 3355–3363, 2017.

[6] Y. Joly, I. N. Feze, L. Song, and B. M. Knoppers, “Comparative approaches to genetic discrimination: chasing shadows?” Trends in Genetics, vol. 33, no. 5, pp. 299–302, 2017.

[7] Z. Wang, M. Gerstein, and M. Snyder, “Rna-seq: a revolutionary tool for transcriptomics,” Nature reviews genetics, vol. 10, no. 1, pp. 57–63, 2009.

[8] F. Tang, C. Barbacioru, Y. Wang, E. Nordman, C. Lee, N. Xu, X. Wang, J. Bodeau, B. B. Tuch, A. Siddiqui et al., “mrna-seq whole-transcriptome analysis of a single cell,” Nature methods, vol. 6, no. 5, pp. 377–382, 2009.

[9] V. Marx, “Method of the year: spatially resolved transcriptomics,” Nature Methods, vol. 18, no. 1, pp. 9–14, 2021.

[10] J. Zou, M. Huss, A. Abid, P. Mohammadi, A. Torkamani, and A. Telenti, “A primer on deep learning in genomics,” Nature genetics, vol. 51, no. 1, pp. 12–18, 2019.

[11] G. A. Kaissis, M. R. Makowski, D. Rückert, and R. F. Braren, “Secure, privacy-preserving and federated machine learning in medical imaging,” Nature Machine Intelligence, vol. 2, no. 6, pp. 305–311, 2020.

[12] A. Esteva, A. Robicquet, B. Ramsundar, V. Kuleshov, M. DePristo, K. Chou, C. Cui, G. Corrado, S. Thrun, and J. Dean, “A guide to deep learning in healthcare,” Nature medicine, vol. 25, no. 1, pp. 24–29, 2019.

[13] M. Al-Rubaie and J. M. Chang, “Privacy-preserving machine learning: Threats and solutions,” IEEE Security & Privacy, vol. 17, no. 2, pp. 49–58, 2019.

[14] M. J. Sheller, B. Edwards, G. A. Reina, J. Martin, S. Pati, A. Kotrotsou, M. Milchenko, W. Xu, D. Marcus, R. R. Colen et al., “Federated learning in medicine: facilitating multi-institutional collaborations without sharing patient data,” Scientific reports, vol. 10, no. 1, pp. 1–12, 2020.

[15] C. G. Schwarz, W. K. Kremers, T. M. Therneau, R. R. Sharp, J. L. Gunter, P. Vemuri, A. Arani, A. J. Spychalla, K. Kantarci, D. S. Knopman et al., “Identification of anonymous mri research participants with face-recognition software,” New England Journal of Medicine, vol. 381, no. 17, pp. 1684–1686, 2019.

[16] A. Harmanci and M. Gerstein, “Quantification of private information leakage from phenotype-genotype data: linking attacks,” Nature methods, vol. 13, no. 3, pp. 251–256, 2016.

[17] B. McMahan, E. Moore, D. Ramage, S. Hampson, and B. A. y Arcas, “Communication-efficient learning of deep networks from decentralized data,” in Artificial intelligence and statistics. PMLR, 2017, pp. 1273–1282.

[18] B. Hitaj, G. Ateniese, and F. Perez-Cruz, “Deep models under the gan: information leakage from collaborative deep learning,” in Proceedings of the 2017 ACM SIGSAC conference on computer and communications security, 2017, pp. 603–618.

[19] L. Melis, C. Song, E. De Cristofaro, and V. Shmatikov, “Exploiting unintended feature leakage in collaborative learning,” in 2019 IEEE Symposium on Security and Privacy (SP). IEEE, 2019, pp. 691–706.

[20] M. Nasr, R. Shokri, and A. Houmansadr, “Comprehensive privacy analysis of deep learning: Passive and active white-box inference attacks against centralized and federated learning,” in 2019 IEEE symposium on security and privacy (SP). IEEE, 2019, pp. 739–753.

[21] L. Zhu, Z. Liu, and S. Han, “Deep leakage from gradients,” Advances in Neural Information Processing Systems, vol. 32, 2019.

[22] V. Tolpegin, S. Truex, M. E. Gursoy, and L. Liu, “Data poisoning attacks against federated learning systems,” in European Symposium on Research in Computer Security. Springer, 2020, pp. 480–501.

[23] M. A. Rahman, T. Rahman, R. Laganiere, N. Mohammed, and Y. Wang, “Membership inference attack against differentially private deep learning model.” Trans. Data Priv., vol. 11, no. 1, pp. 61–79, 2018.

[24] H. Lu, C. Liu, T. He, S. Wang, and K. S. Chan, “Sharing models or coresets: A study based on membership inference attack,” arXiv preprint 2007.02977, 2020.

[25] A. Salem, Y. Zhang, M. Humbert, P. Berrang, M. Fritz, and M. Backes, “Ml-leaks: Model and data independent membership inference attacks and defenses on machine learning models,” arXiv preprint 1806.01246, 2018.

[26] H. Hu, Z. Salcic, L. Sun, G. Dobbie, and X. Zhang, “Source inference attacks in federated learning,” arXiv preprint 2109.05659, 2021.

[27] J. Geiping, H. Bauermeister, H. DrÖge, and M. Moeller, “Inverting gradients – how easy is it to break privacy in federated learning?” 2020.

[28] H. J. La, M. K. Kim, and S. D. Kim, “A personal healthcare system with inference-as-a-service,” in 2015 IEEE International Conference on Services Computing. IEEE, 2015, pp. 249–255.

[29] K. Bonawitz, V. Ivanov, B. Kreuter, A. Marcedone, H. B. McMahan, S. Patel, D. Ramage, A. Segal, and K. Seth, “Practical secure aggregation for privacy-preserving machine learning,” in proceedings of the 2017 ACM SIGSAC Conference on Computer and Communications Security, 2017, pp. 1175–1191.

[30] S. Wagh, D. Gupta, and N. Chandran, “Securenn: 3-party secure computation for neural network training.” Proc. Priv. Enhancing Technol., vol. 2019, no. 3, pp. 26–49, 2019.

[31] S. Sav, A. Pyrgelis, J. R. Troncoso-Pastoriza, D. Froelicher, J.-P. Bossuat, J. S. Sousa, and J.-P. Hubaux, “Poseidon: privacy-preserving federated neural network learning,” arXiv preprint 2009.00349, 2020.

[32] J. Zhou, L. Zhou, D. Wang, X. Xu, H. Li, Y. Chu, W. Han, and X. Gao, “Personalized and privacy-preserving federated heterogeneous medical image analysis with pppml-hmi,” medRxiv, pp. 2023–02, 2023.

[33] D. Froelicher, J. R. Troncoso-Pastoriza, J. L. Raisaro, M. A. Cuendet, J. S. Sousa, H. Cho, B. Berger, J. Fellay, and J.-P. Hubaux, “Truly privacy-preserving federated analytics for precision medicine with multiparty homomorphic encryption,” Nature communications, vol. 12, no. 1, pp. 1–10, 2021.

[34] S. Warnat-Herresthal, H. Schultze, K. L. Shastry, S. Manamohan, S. Mukherjee, V. Garg, R. Sarveswara, K. Händler, P. Pickkers, N. A. Aziz et al., “Swarm learning for decentralized and confidential clinical machine learning,” Nature, vol. 594, no. 7862, pp. 265–270, 2021.

[35] M. Ali, H. Karimipour, and M. Tariq, “Integration of blockchain and federated learning for internet of things: Recent advances and future challenges,” Computers & Security, vol. 108, p. 102355, 2021.

[36] C. Dwork, “Differential privacy: A survey of results,” in International conference on theory and applications of models of computation. Springer, 2008, pp. 1–19.

[37] K. Wei, J. Li, M. Ding, C. Ma, H. H. Yang, F. Farokhi, S. Jin, T. Q. Quek, and H. V. Poor, “Federated learning with differential privacy: Algorithms and performance analysis,” IEEE Transactions on Information Forensics and Security, vol. 15, pp. 3454–3469, 2020.

[38] N. Rodríguez-Barroso, G. Stipcich, D. Jiménez-López, J. A. Ruiz-Millán, E. Martínez-Cámara, G. González-Seco, M. V. Luzón, M. A. Veganzones, and F. Herrera, “Federated learning and differential privacy: Software tools analysis, the sherpa. ai fl framework and methodological guidelines for preserving data privacy,” Information Fusion, vol. 64, pp. 270–292, 2020.

[39] R. Liu, Y. Cao, H. Chen, R. Guo, and M. Yoshikawa, “Flame: Differentially private federated learning in the shuffle model,” arXiv preprint 2009.08063, 2020.

[40] A. Girgis, D. Data, S. Diggavi, P. Kairouz, and A. T. Suresh, “Shuffled model of differential privacy in federated learning,” in International Conference on Artificial Intelligence and Statistics. PMLR, 2021, pp. 2521–2529.

[41] B. Ghazi, R. Kumar, P. Manurangsi, R. Pagh, and A. Sinha, “Differentially private aggregation in the shuffle model: Almost central accuracy in almost a single message,” in International Conference on Machine Learning. PMLR, 2021, pp. 3692–3701.

[42] S. D. Constable, Y. Tang, S. Wang, X. Jiang, and S. Chapin, “Privacy-preserving gwas analysis on federated genomic datasets,” in BMC medical informatics and decision making, vol. 15, no. 5. BioMed Central, 2015, pp. 1–9.

[43] H. Cho, D. J. Wu, and B. Berger, “Secure genome-wide association analysis using multiparty computation,” Nature biotechnology, vol. 36, no. 6, pp. 547–551, 2018.

[44] G. Gü rsoy, E. Chielle, C. M. Brannon, M. Maniatakos, and M. Gerstein, “Privacy-preserving genotype imputation with fully homomorphic encryption,” Cell Systems, vol. 13, no. 2, pp. 173–182, 2022.

[45] Z. He, Y. Li, J. Li, K. Li, Q. Cai, and Y. Liang, “Achieving differential privacy of genomic data releasing via belief propagation,” Tsinghua Science and Technology, vol. 23, no. 4, pp. 389–395, 2018.

[46] E. Yilmaz, T. Ji, E. Ayday, and P. Li, “Genomic data sharing under dependent local differential privacy,” arXiv preprint 2102.07357, 2021.

[47] N. Almadhoun, E. Ayday, and Ö. Ulusoy, “Differential privacy under dependent tuples—the case of genomic privacy,” Bioinformatics, vol. 36, no. 6, pp. 1696–1703, 2020.

[48] G. Gü rsoy, C. M. Brannon, F. C. Navarro, and M. Gerstein, “Fancy: fast estimation of privacy risk in functional genomics data,” Bioinformatics, vol. 36, no. 21, pp. 5145–5150, 2020.

[49] G. Gü rsoy, T. Li, S. Liu, E. Ni, C. M. Brannon, and M. B. Gerstein, “Functional genomics data: privacy risk assessment and technological mitigation,” Nature Reviews Genetics, pp. 1–14, 2021.

[50] O. Zolotareva, R. Nasirigerdeh, J. Matschinske, R. Torkzadehmahani, M. Bakhtiari, T. Frisch, J. Späth, D. B. Blumenthal, A. Abbasinejad, P. Tieri et al., “Flimma: a federated and privacy-aware tool for differential gene expression analysis,” Genome biology, vol. 22, no. 1, pp. 1–26, 2021.

[51] C. Beguier, J. O. d. Terrail, I. Meah, M. Andreux, and E. W. Tramel, “Differentially private federated learning for cancer prediction,” arXiv preprint 2101.02997, 2021.

[52] W. Li, F. Milletarì, D. Xu, N. Rieke, J. Hancox, W. Zhu, M. Baust, Y. Cheng, S. Ourselin, M. J. Cardoso et al., “Privacy-preserving federated brain tumour segmentation,” in International workshop on machine learning in medical imaging. Springer, 2019, pp. 133–141.

[53] G. Kaissis, A. Ziller, J. Passerat-Palmbach, T. Ryffel, D. Usynin, A. Trask, I. Lima, J. Mancuso, F. Jungmann, M.-M. Steinborn et al., “End-to-end privacy preserving deep learning on multiinstitutional medical imaging,” Nature Machine Intelligence, vol. 3, no. 6, pp. 473–484, 2021.

[54] E. Shapiro, T. Biezuner, and S. Linnarsson, “Single-cell sequencing-based technologies will revolutionize whole-organism science,” Nature Reviews Genetics, vol. 14, no. 9, pp. 618–630, 2013.

[55] D. J. Burgess, “Spatial transcriptomics coming of age,” Nature Reviews Genetics, vol. 20, no. 6, pp. 317–317, 2019.

[56] B. He, L. Bergenstråhle, L. Stenbeck, A. Abid, A. Andersson, å. Borg, J. Maaskola, J. Lundeberg, and J. Zou, “Integrating spatial gene expression and breast tumour morphology via deep learning,” Nature biomedical engineering, vol. 4, no. 8, pp. 827–834, 2020.

[57] R. A. Fisher and F. Yates, Statistical tables for biological, agricultural and medical research. Hafner Publishing Company, 1953.

[58] T. Li and N. Li, “On the tradeoff between privacy and utility in data publishing,” in Proceedings of the 15th ACM SIGKDD international conference on Knowledge discovery and data mining, 2009, pp. 517–526.

[59] U. C. Bureau, “Disclosure avoidance for the 2020 census: An introduction,” Nov 2021. [Online]. Available: https://www.census.gov/library/publications/2021/decennial/2020-census-disclosure-avoidance-handbook.html

[60] V. Feldman, I. Mironov, K. Talwar, and A. Thakurta, “Privacy amplification by iteration,” in 2018 IEEE 59th Annual Symposium on Foundations of Computer Science (FOCS). IEEE, 2018, pp. 521–532.

[61] J. H. Cheon, A. Kim, M. Kim, and Y. Song, “Homomorphic encryption for arithmetic of approximate numbers,” in International Conference on the Theory and Application of Cryptology and Information Security. Springer, 2017, pp. 409–437.

[62] D. Tran, H. Nguyen, B. Tran, C. La Vecchia, H. N. Luu, and T. Nguyen, “Fast and precise single-cell data analysis using a hierarchical autoencoder,” Nature communications, vol. 12, no. 1, pp. 1–10, 2021.

[63] S. Cristea and K. Polyak, “Dissecting the mammary gland one cell at a time,” Nature communications, vol. 9, no. 1, pp. 1–3, 2018.

[64] L. González-Silva, L. Quevedo, and I. Varela, “Tumor functional heterogeneity unraveled by scrna-seq technologies,” Trends in cancer, vol. 6, no. 1, pp. 13–19, 2020.

[65] Q. Zhang, J. Wang, P. Wang, T. Tang, P. Li, Y. Pei, X. Zhang, W. Zhang, Q. Gu, and Q. Ji, “Establishment and optimization of scrna-seq assay to find the mechanism of immune therapy against tumors,” Cell, vol. 8, p. 9, 2021.

[66] S. Ding, X. Chen, and K. Shen, “Single-cell rna sequencing in breast cancer: Understanding tumor heterogeneity and paving roads to individualized therapy,” Cancer Communications, vol. 40, no. 8, pp. 329–344, 2020.

[67] A. Butler, P. Hoffman, P. Smibert, E. Papalexi, and R. Satija, “Integrating single-cell transcriptomic data across different conditions, technologies, and species,” Nature biotechnology, vol. 36, no. 5, pp. 411–420, 2018.

[68] V. Y. Kiselev, K. Kirschner, M. T. Schaub, T. Andrews, A. Yiu, T. Chandra, K. N. Natarajan, W. Reik, M. Barahona, A. R. Green et al., “Sc3: consensus clustering of single-cell rna-seq data,” Nature methods, vol. 14, no. 5, pp. 483–486, 2017.

[69] P. Lin, M. Troup, and J. W. Ho, “Cidr: Ultrafast and accurate clustering through imputation for single-cell rna-seq data,” Genome biology, vol. 18, no. 1, pp. 1–11, 2017.

[70] M. Guo, H. Wang, S. S. Potter, J. A. Whitsett, and Y. Xu, “Sincera: a pipeline for single-cell rna-seq profiling analysis,” PLoS computational biology, vol. 11, no. 11, p. e1004575, 2015.

[71] L. Zhang, Z. Li, K. M. Skrzypczynska, Q. Fang, W. Zhang, S. A. O’Brien, Y. He, L. Wang, Q. Zhang, A. Kim et al., “Single-cell analyses inform mechanisms of myeloid-targeted therapies in colon cancer,” Cell, vol. 181, no. 2, pp. 442–459, 2020.

[72] C. Song, T. Ristenpart, and V. Shmatikov, “Machine learning models that remember too much,” in Proceedings of the 2017 ACM SIGSAC Conference on computer and communications security, 2017, pp. 587–601.

[73] A. Raj, Y. Bresler, and B. Li, “Improving robustness of deeplearning-based image reconstruction,” in International Conference on Machine Learning. PMLR, 2020, pp. 7932–7942.

[74] R. Cappelli, D. Maio, A. Lumini, and D. Maltoni, “Fingerprint image reconstruction from standard templates,” IEEE transactions on pattern analysis and machine intelligence, vol. 29, no. 9, pp. 1489–1503, 2007.

[75] B. Zhao, K. R. Mopuri, and H. Bilen, “idlg: Improved deep leakage from gradients,” arXiv preprint 2001.02610, 2020.

[76] B. Balle and Y.-X. Wang, “Improving the gaussian mechanism for differential privacy: Analytical calibration and optimal denoising,” in International Conference on Machine Learning. PMLR, 2018, pp. 394–403.

[77] Ú. Erlingsson, V. Feldman, I. Mironov, A. Raghunathan, K. Talwar, and A. Thakurta, “Amplification by shuffling: From local to central differential privacy via anonymity,” in Proceedings of the Thirtieth Annual ACM-SIAM Symposium on Discrete Algorithms. SIAM, 2019, pp. 2468–2479.

[78] L. Yan, M. Yang, H. Guo, L. Yang, J. Wu, R. Li, P. Liu, Y. Lian, X. Zheng, J. Yan et al., “Single-cell rna-seq profiling of human preimplantation embryos and embryonic stem cells,” Nature structural & molecular biology, vol. 20, no. 9, pp. 1131–1139, 2013.

[79] A. A. Pollen, T. J. Nowakowski, J. Shuga, X. Wang, A. A. Leyrat, J. H. Lui, N. Li, L. Szpankowski, B. Fowler, P. Chen et al., “Lowcoverage single-cell mrna sequencing reveals cellular heterogeneity and activated signaling pathways in developing cerebral cortex,” Nature biotechnology, vol. 32, no. 10, pp. 1053–1058, 2014.

[80] S. Hrvatin, D. R. Hochbaum, M. A. Nagy, M. Cicconet, K. Robertson, L. Cheadle, R. Zilionis, A. Ratner, R. Borges-Monroy, A. M. Klein et al., “Single-cell analysis of experience-dependent transcriptomic states in the mouse visual cortex,” Nature neuroscience, vol. 21, no. 1, pp. 120–129, 2018.

[81] A. Paszke, S. Gross, F. Massa, A. Lerer, J. Bradbury, G. Chanan, T. Killeen, Z. Lin, N. Gimelshein, L. Antiga et al., “Pytorch: An imperative style, high-performance deep learning library,” Advances in neural information processing systems, vol. 32, pp. 8026–8037, 2019.

[82] B. Jayaraman and D. Evans, “Evaluating differentially private machine learning in practice,” in 28th USENIX Security Symposium (USENIX Security 19), 2019, pp. 1895–1912.

[83] M. Fredrikson, E. Lantz, S. Jha, S. Lin, D. Page, and T. Ristenpart, “Privacy in pharmacogenetics: An end-to-end case study of personalized warfarin dosing,” in 23rd {USENIX} Security Symposium ({USENIX} Security 14), 2014, pp. 17–32.

[84] M. Fredrikson, S. Jha, and T. Ristenpart, “Model inversion attacks that exploit confidence information and basic countermeasures,” in Proceedings of the 22nd ACM SIGSAC Conference on Computer and Communications Security, ser. CCS ‘15. New York, NY, USA: Association for Computing Machinery, 2015, p. 1322–1333. [Online]. Available: https://doi.org/10.1145/2810103.2813677

[85] A. Benaissa, B. Retiat, B. Cebere, and A. E. Belfedhal, “Tenseal: A library for encrypted tensor operations using homomorphic encryption,” arXiv preprint 2104.03152, 2021.

